# Thermodynamic integration for dynamic causal models

**DOI:** 10.1101/471417

**Authors:** Eduardo A. Aponte, Sudhir Raman, Stefan Frässle, Jakob Heinzle, Will D. Penny, Klaas E. Stephan

## Abstract

In generative modeling of neuroimaging data, such as dynamic causal modeling (DCM), one typically considers several alternative models, either to determine the most plausible explanation for observed data (Bayesian model selection) or to account for model uncertainty (Bayesian model averaging). Both procedures rest on estimates of the model evidence, a principled trade-off between model accuracy and complexity. In DCM, the log evidence is usually approximated using variational Bayes (VB) under the Laplace approximation (VBL). Although this approach is highly efficient, it makes distributional assumptions and can be vulnerable to local extrema. An alternative to VBL is Markov Chain Monte Carlo (MCMC) sampling, which is asymptotically exact but orders of magnitude slower than VB. This has so far prevented its routine use for DCM.

This paper makes four contributions. First, we introduce a powerful MCMC scheme – thermodynamic integration (TI) – to neuroimaging and present a derivation that establishes a theoretical link to VB. Second, this derivation is based on a tutorial-like introduction to concepts of free energy in physics and statistics. Third, we present an implementation of TI for DCM that rests on population MCMC. Fourth, using simulations and empirical functional magnetic resonance imaging (fMRI) data, we compare log evidence estimates obtained by TI, VBL, and other MCMC-based estimators (prior arithmetic mean and posterior harmonic mean). We find that model comparison based on VBL gives reliable results in most cases, justifying its use in standard DCM for fMRI. Furthermore, we demonstrate that for complex and/or nonlinear models, TI may provide more robust estimates of the log evidence. Importantly, accurate estimates of the model evidence can be obtained with TI in acceptable computation time. This paves the way for using DCM in scenarios where the robustness of single-subject inference and model selection becomes paramount, such as differential diagnosis in clinical applications.

## Introduction

Dynamic causal models (DCMs; Friston et al., 2003) are generative models that serve to infer latent neurophysiological processes and circuit properties – e.g., the effective connectivity between neuronal populations – from neuroimaging measurements such as functional magnetic resonance imaging (fMRI) or magneto-/electroencephalography (M/EEG) data. As reviewed by (Daunizeau et al., 2011), DCMs consist of two hierarchically related layers: a set of state equations describing neuronal population activity, and an observation model which links neurophysiological states to observed signals and accounts for measurement noise. Equipped with a prior distribution over model parameters, a DCM specifies a full generative or probabilistic forward model that can be inverted using Bayesian techniques.

Depending on the specific type of DCM, interactions between neuronal populations and the generation of measurable signals – e.g., blood oxygen level dependent (BOLD) signal in fMRI or scalp voltage fluctuations in EEG – are represented by different nonlinear equations. These nonlinearities prevent exact analytical inference and require the use of approximate or asymptotic inference techniques. To date, variational Bayes under the Laplace approximation (VBL; Friston et al., 2007) has been the method of choice for DCM, partially because of its computational efficiency.

In addition to inference on model parameters, an important scientific problem is the comparison of competing hypotheses that are formalized as different models. Under the Bayesian framework, model comparison is based on the evidence or marginal likelihood of a model. The evidence corresponds to the denominator or normalization constant from Bayes’ theorem and represents the probability of the observed data under a given model. It is a widely used score of model quality that quantifies the trade-off between model fit and complexity (Bishop, 2006; MacKay, 2002).

Unfortunately, in most instances, it is not feasible to derive an analytical expression of the model evidence due to the intractable integrals that arise from the marginalization of the model parameters. Various asymptotical approximations exist, such as the Bayesian Information Criterion (BIC; Schwarz, 1978) and more recently the Widely Applicable Bayesian Information Criterion (WBIC; Watanabe, 2013). Within the framework of variational Bayes (VB), a lower bound approximation of the log model evidence (LME) is obtained as a byproduct of model inversion: the variational negative free energy (which we refer to as −*F_VB_* throughout this paper).

While highly efficient, model comparison based on the variational free energy has several potential pitfalls. For example, under the Laplace approximation as used in the context of DCM, there is no guarantee that −*F_VB_* still represents a lower bound of the LME (Wipf and Nagarajan, 2009). Furthermore, VB is commonly performed in combination with a mean field approximation, and the effect of this approximation on the posterior estimates can be difficult to predict (for discussion, see Daunizeau et al., 2011). Finally, in non-linear models, the posterior could become a multimodal density, a condition that aggravates the application of gradient ascent methods regularly used in combination with the Laplace approximation.

For these reasons, Markov Chain Monte Carlo (MCMC) sampling has been explored as an alternative inference technique for DCM (Aponte et al., 2016; Chumbley et al., 2007; Penny and Sengupta, 2016; Raman et al., 2016; Sengupta et al., 2016; 2015). MCMC is particularly attractive for variants of DCMs in which Gaussian assumptions might be less adequate, such as nonlinear DCMs for fMRI (Stephan et al., 2008), DCMs of electrophysiological data (Moran et al., 2013a), or DCMs for layered fMRI signals (Heinzle et al., 2016). MCMC is also useful when extending DCM to more complex hierarchical models (Raman et al., 2016), in which the derivation of update equations for VB becomes difficult (but see Yao et al., 2018). MCMC does not entail any assumptions about the posterior distribution and is asymptotically exact. However, in practice, its computational cost leads to runtimes that are often prohibitively long for the datasets and models commonly encountered in neuroimaging. Furthermore, in contrast to VB, MCMC-based inversion of generative models does not provide an estimate of the model evidence for free.

While several MCMC strategies for computing the model evidence in neuroimaging applications have been explored (e.g., Aponte et al., 2016; Penny and Sengupta, 2016; Raman et al., 2016), one particularly powerful and theoretically attractive MCMC variant that has not yet been investigated in detail is thermodynamic integration (TI) (but see Penny and Sengupta, 2016). This method, like VB, rests on the concept of free energy and has been proposed as gold standard for model evidence estimation (Calderhead and Girolami, 2009; Lartillot and Philippe, 2006). Despite strong theoretical advantages, so far, the computational costs of TI have prohibited its practical use in neuroimaging.

This technical note introduces TI to neuroimaging in general and DCM in particular, with three distinct contributions. First, we present the theoretical foundations of TI and demonstrate its theoretical link to VB. Second, we present an efficient implementation of TI that rests on population MCMC and parallelization using graphical processing units (GPUs). Third, we evaluate our TI scheme using simulations and empirical fMRI data. Specifically, we compare LME estimates obtained by TI to those by conventional sampling-based estimates of the LME (prior arithmetic mean and posterior harmonic mean) and VBL.

This paper is organized as follows: We begin with a brief review of DCM for fMRI to keep the paper self-contained. We then turn to model comparison, first reviewing conventional sampling-based estimators of the LME and subsequently presenting a derivation of TI that reveals its theoretical relation to VB. Using simulated data, we verify the accuracy of our TI implementation and its superiority compared to conventional sampling-based estimators of the LME. Finally, we compare the ranking of competing DCMs by TI and VBL for two empirical fMRI datasets – one frequently used dataset on visual attention (Buchel and Friston, 1997) with nonlinear DCMs that may pose a particular challenge for VBL, and a more recent dataset on face perception (Frassle et al., 2016a).

## Methods

### Dynamic Causal Models

This paper focuses on fMRI data, and we therefore limit our discussion of DCM to BOLD signals (Friston et al., 2003; Stephan et al., 2008; 2007). In brief, DCM for fMRI is characterized by two layers: first, a set of ordinary differential equations that model the dynamics of interacting neuronal states *x* and local hemodynamic states *h*. Second, the hemodynamic states enter a static nonlinear observation equation that relates venous blood volume and deoxyhemoglobin content to measured BOLD signal changes. In the following, we discuss only the most relevant equations, in order to convey an understanding of the type of problem that model inversion in DCM faces.

The general form of the dynamics of the neuronal layer is

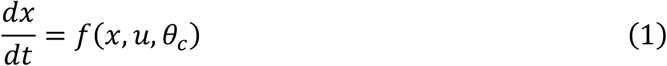

where *x* = (*x*_2_,…,*x_N_*) describes the neuronal states of *N* regions, *u* = (*u*_2_,…,*u_M_*) represents the time series of *M* experimental manipulations or inputs, and *θ_c_* are the connectivity parameters that determine the neuronal dynamics. Using a second order Taylor expansion (Stephan et al., 2008), the dynamics *f* can be approximated as:

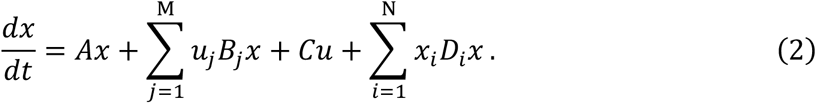

The connectivity parameters *θ_c_* can be divided into four subsets: The *N* × *N* matrix *A* describes endogenous connectivity strengths between regions. The set of *N* × *N* matrices *B* = {*B*_1_,…,*B_M_*} encodes modulatory effects of inputs on connections between regions. The *N* × *M* matrix *C* describes the direct effects of driving inputs on regions. Finally, the *N* × *N* matrices *D* = {*D*_1_,…,*D_N_*} denote second-order interactions between two regions that affect a third one. Linear DCMs use *A* and *C* matrices, bilinear DCMs contain at least one non-zero *B* matrix, and nonlinear DCMs contain at least one non-zero *D* matrix. Together *θ_c_* = {*A, B, C, D*} fully describe the dynamics of the neuronal layer.

The hemodynamic model of DCM originates from the Balloon model proposed by Buxton et al. (1998) and extended by Friston et al. (2000) and Stephan et al. (2007). In brief, it describes how changes in neuronal states locally alter cerebral blood flow, which, in turn, affects venous blood volume and deoxyhemoglobin content. The model consists of a cascade of deterministic differential equations:

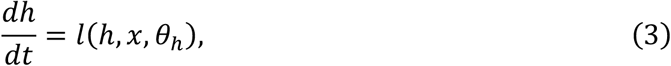

where *h* = (*h*_1_,…,*h_N_*) denotes hemodynamic states in each of *N* regions. Detailed equations and the meaning of the hemodynamic parameters *θ_h_* can be found in Stephan et al. (2007). It is worth noting that the hemodynamic equations are nonlinear and that the original implementation in SPM uses a local (bi)linear approximation (Friston et al., 2003).

Finally, hemodynamic states enter a static nonlinear observation equation *g* with parameters *θ_g_* that models the BOLD signal *y*:

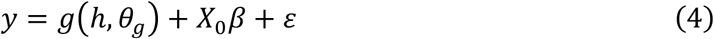

The term *X*_0_ is a matrix of confound regressors that accounts for constant terms and low frequency fluctuations. The Gaussian observation noise *ε* is characterized by the covariance matrix:

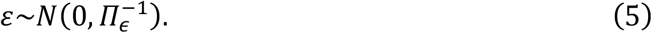

The precision matrix *Π_ϵ_* is represented as a linear combination *Π_ϵ_* = Σ_*r*_ exp(*λ_r_*)*Q_r_*. The precision components *Q_r_* serve to account for temporal autocorrelation and regional differences in noise variance (Friston et al., 2003). Here, we assume that the time series have been whitened and therefore only account for region-specific variances. In this case, each *Q_r_* is a diagonal matrix with diagonal elements belonging to region *r* set to 1, and 0 elsewhere.

To complete the generative model, the prior distribution of the parameters *Θ* = (*θ_c_,θ_h_,θ_g_, β*) and hyperparameters *Λ* needs to be specified. Here, the priors have been largely matched to SPM8 release 5236 (http://www.fil.ion.ucl.ac.uk/spm), except for the scaling of the prior variance of the coefficients of the confound matrix *X*_0_, which was adapted to the scaling of the data as explained in Supp. 1. All parameters’ prior distributions are Gaussian, and when positivity needs to be enforced, an adequate transformation function is used. A detailed specification of the priors is provided by Sup. Table 1.

### Bayesian model comparison and selection

Bayesian inference involves the specification of a probabilistic or generative model *m* with data *y* and parameters *θ*. The model has two components: the prior density over *θ, p*(*θ|m*), and the likelihood function *p*(*y|θ,m*). These are combined to form the posterior distribution using Bayes’ theorem. Conditioning on a given model *m*, the posterior distribution is:

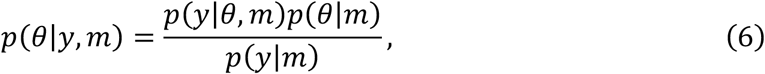

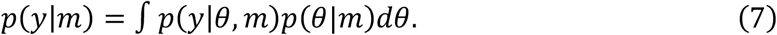

The normalization constant in the denominator, *p*(*y|m*), is known as the marginal likelihood or model evidence and corresponds to the likelihood of the data after marginalizing out the parameters of the model.

In practice, given the monotonicity of the logarithmic function, either the evidence or its logarithm can be used to score a set of candidate models *m*_1_,…,*m_n_* (Bayesian model comparison) and to identify the best model within the model space studied (Bayesian model selection; BMS). One common metric for assessing the relative goodness of two models is the Bayes factor (Kass and Raftery, 1995):

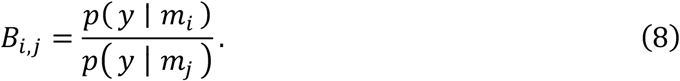

or, equivalently, the exponential of the difference in LME of two models.

BMS has gained an important role in neuroimaging, not only for DCM but also in other contexts requiring model comparison, such as EEG source reconstruction (Henson et al., 2009; Wipf and Nagarajan, 2009), or computational neuroimaging (Friston and Dolan, 2010; Stephan et al., 2015; 2017a). Group-level BMS techniques exist which account for individual heterogeneity by treating the model as a random variable in the population (Friston et al., 2016; Rigoux et al., 2014; Stephan et al., 2009). Finally, Bayesian model averaging allows one to compute an average posterior over models (Penny et al., 2010; Trujillo-Barreto et al., 2004), weighted by the posterior probability of each model. Critically, these approaches rely on an accurate estimate of each model’s evidence.

As mentioned above, except for some special cases, the model evidence cannot be determined analytically, and one typically has to resort to approximations. One computationally efficient option is VB (for textbook treatments, see Koller and Friedman, 2009; MacKay, 2003), which provides a lower bound of the LME. An alternative, which we explore in detail here, is MCMC sampling. This family of methods is characterized by simulating a Markov process whose stationary distribution corresponds to the posterior distribution *p*(*θ|y, m*) (for a textbook reference, see Robert and Casella, 2013).

In the next sections, we describe standard sampling-based estimators of the model evidence and juxtapose these to the variational negative free energy approximation in VBL. We do not consider classical approximations to the LME such as BIC, since these have already been evaluated in the context of DCM in previous work and were found to be less adequate than the variational negative free energy (Penny, 2012).

### Prior arithmetic mean estimator (AME)

Importance sampling is a Monte Carlo method for approximating the expected value of a random variable *h*(*X*) under the density *p* by means of an auxiliary density function *w*(*X*), which is required to be absolutely continuous with respect to *p* (Robert and Casella, 2013; p. 92, Def. 3.9), or less formally, the auxiliary density *w* should share the same support as *p* to avoid zeros in the denominator:

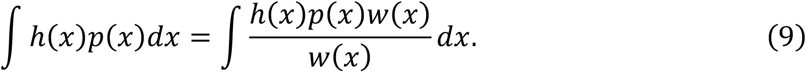

From the strong law of large numbers, if this expected value exists, the process

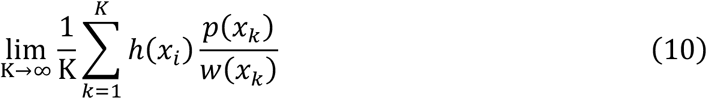

converges almost surely to Eq. 9 when the samples *x*_1_,…,*x_K_* have been drawn from the auxiliary distribution *w*.

In order to approximate the model evidence by importance sampling, the simplest choice of the auxiliary density is the prior distribution, *w* = *p*(*θ | m*). This results in the prior arithmetic mean estimator (AME):

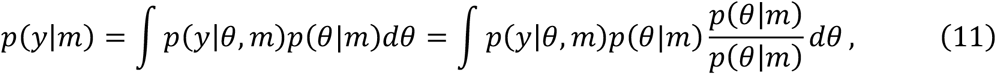

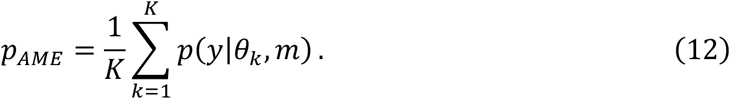

where samples *θ_k_* have been obtained from the prior distribution *p*(*θ|m*). Because samples of the likelihood *p*(*y|θ,m*) can greatly exceed the range of double precision floating point numbers, it is necessary to normalize the likelihood function in log space. This can be achieved with the following formula:

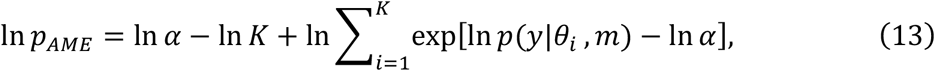

where *α* > 0 is an arbitrary constant. In all analyses reported here, *α* was set to 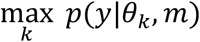.

A serious shortcoming of AME is that in the great majority of situations most samples drawn from the prior have very low likelihood. Therefore, an extremely large number of samples is required to ensure that high likelihood regions of the parameter space are taken into account by the estimator; otherwise, the estimator suffers from high variance, as demonstrated previously (Vyshemirsky and Girolami, 2008).

### Posterior harmonic mean estimator (HME)

The second choice for the auxiliary density is the posterior distribution, which results in the posterior harmonic mean estimator (HME). This estimator has received divergent appraisals in the literature as a method for computing the LME (for example Kass and Raftery, 1995; Wolpert and Schmidler, 2012). Re-expressing the model evidence, the HME can be derived as follows:

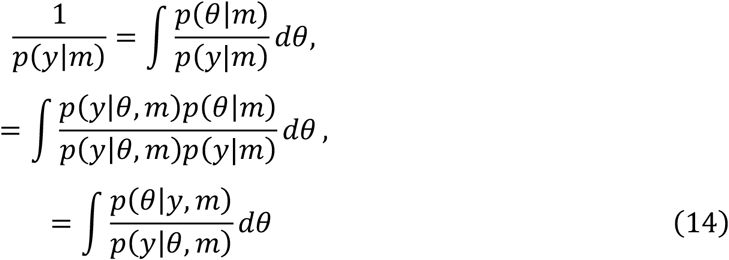

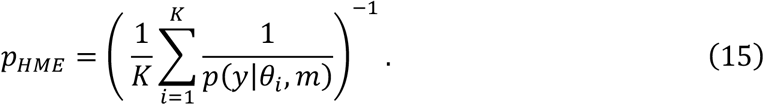

Here, samples *θ_i_* are drawn from the posterior distribution *p*(*θ|y,m*).

In order to avoid numerical instabilities, it is again necessary to normalize in log space, using the formula

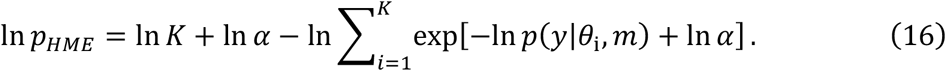

Here, ln *α* has been chosen to be 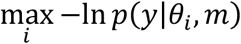.

A disadvantage of HME is that its variance might be infinite when the likelihood function is not heavy-tailed (Raftery et al., 2006), which has serious consequences for the convergence rate of a wide variety of models (Wolpert and Schmidler, 2012). A second problem is that the samples used for HME are obtained from the posterior distribution only. This leads to the opposite behavior as for AME: because the contribution of the prior to the LME might not be appropriately accounted for, the HME tends to overestimate the model evidence, a behavior that can be difficult to diagnose (Lartillot and Philippe, 2006). Several improvements of the HME have been proposed to account for this shortcoming (for example, Raftery et al., 2006).

### Thermodynamic Integration (TI)

This section introduces TI from a statistical physics perspective. Statistical physics is a branch of modern physics that uses methods from probability theory and statistics to characterize the (e.g., thermodynamic) behavior of physical systems. One of the key concepts in statistical physics is that the probability of a particle being in a given state follows a probability density, and that all physically relevant quantities can be derived once this distribution is known. We here focus on the free energy that has a direct relation to the log model evidence in statistics. This also provides a link to the variational Bayes approach conventionally used in DCM to approximate the log evidence (this is described in the subsequent section). Fig. 1 summarizes the analogies in free energy concepts between statistical physics and statistics.

**Figure 1:**
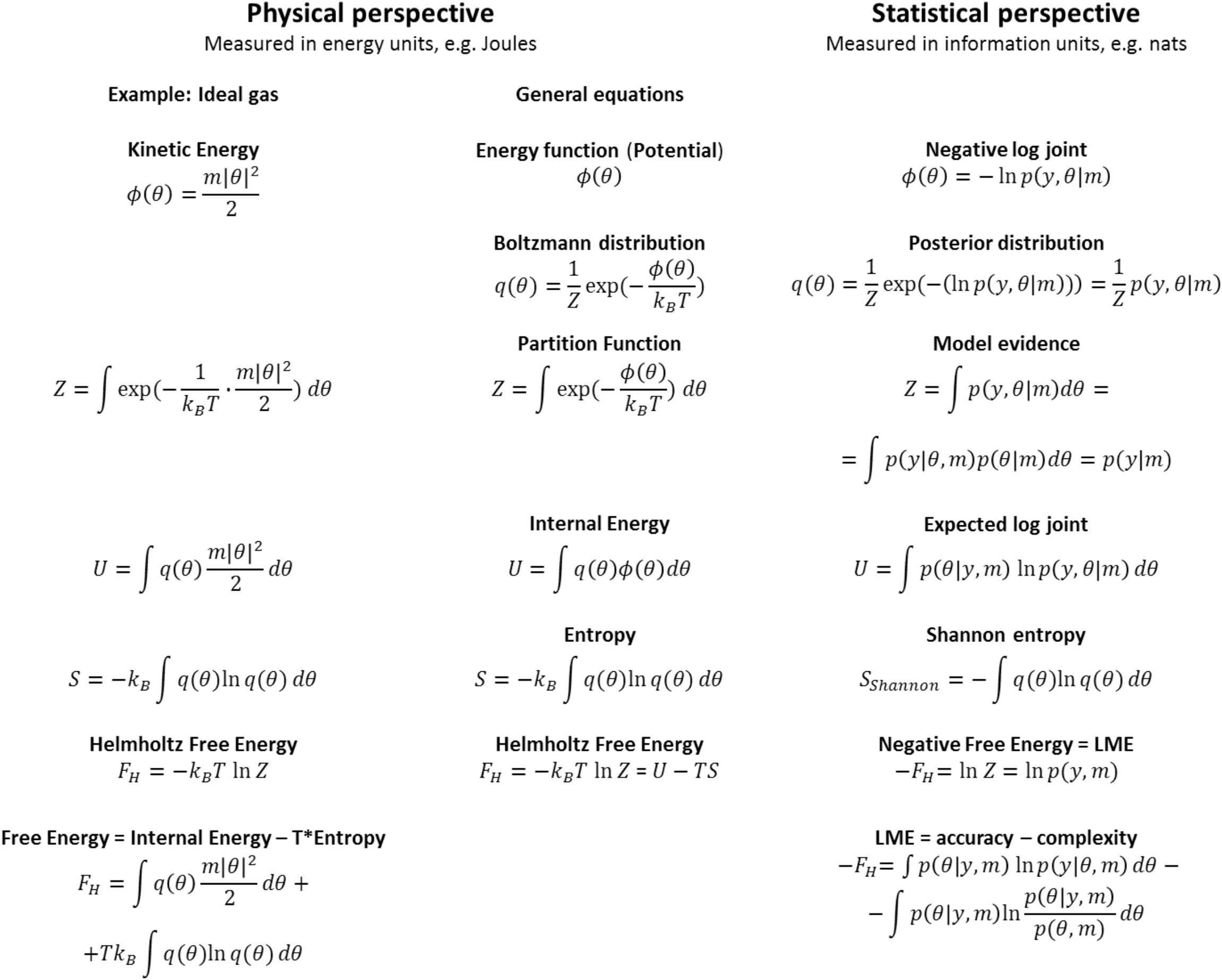
Analogies between concepts of free energy in statistical physics and Bayesian statistics.

#### Free energy: A perspective from physics

In thermodynamics, the analogue of the model evidence (Eq. 7) is the so-called partition function *Z* of a system that consists of an ensemble of particles in thermal equilibrium. A classical discussion of the relationships presented here can be found in Jaynes (1957) and a more modern perspective in (Ortega and Braun, 2013). For example, let us consider an ideal monoatomic gas in which the kinetic energy 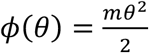 of individual particles is a function of their velocity *θ*. If the system is large enough, the velocity of a single particle can be treated as a continuous random variable. The internal energy *U* of this ideal gas is proportional to the expected energy per particle. It is computed as the weighted sum of the energies *ϕ*(*θ*) associated with all possible velocities, where the weights are given by the probability *q*(*θ*) of the particle being at a certain energy level (i.e., velocity):

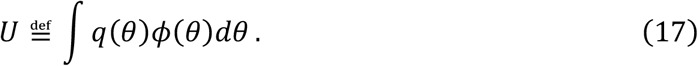

A second important quantity in statistical physics is the entropy *S* of *q*:

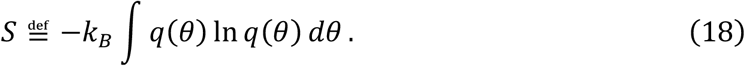

Here, *k_B_* is the Boltzmann constant with units of energy per degree temperature. For an isolated system (i.e., no exchange of matter or energy with the environment), the second law of thermodynamics states that its entropy can only increase or stay constant. Thus, the system is at equilibrium when the associated entropy is maximized, subject to the constraint that the system’s internal energy is constant and equal to *U*, and that *q* is a proper density

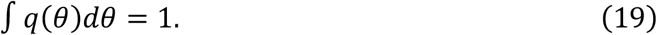

The maximization problem under the above-mentioned conditions can be solved using a variational Lagrangian with two constraints (represented by the Lagrange multipliers *λ*_1_ and *λ*_2_; Blundell and Blundell, 2009; Jaynes, 1957):

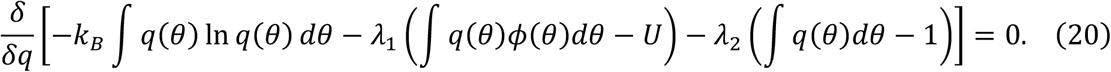

Noting that

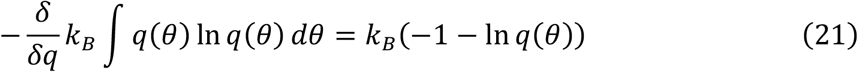

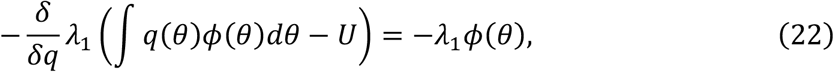

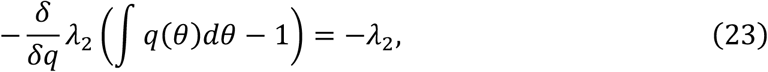

the Lagrangian yields

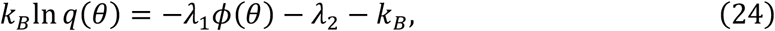

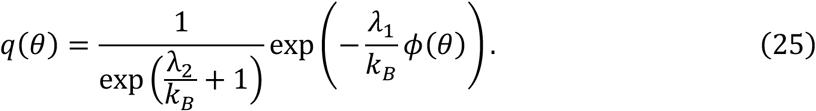

The term *λ*_1_ constitutes the definition of inverse temperature in statistical physics (Blundell and Blundell, 2009; Jaynes, 1957):

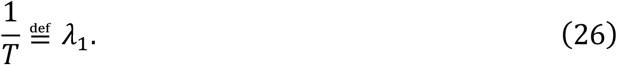

The term 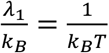 is commonly represented by the symbol *β*. In order to derive the second constant *λ*_2_, we write:

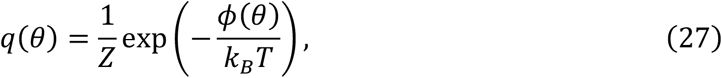

where *Z* is referred to as the partition function of the system:

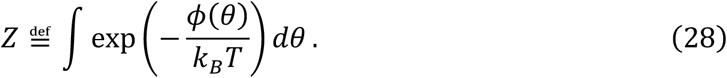

Hence, the term 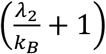 is the normalization constant of *q*(*θ*), and thus *λ*_2_ = *k_B_*(ln *Z* − 1).

With these derivations in mind, we can now approach the physical definition of free energy. In a closed system, the Helmholtz free energy *F_H_* is defined as the difference between the internal energy *U* of the system and its entropy *S* times the temperature *T*:

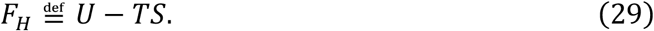

The Helmholtz free energy corresponds to the work (i.e., non-thermal energy in joules that is passed from the system to its environment) that can be attained from a closed system. From Eq. 29, we see that the system with constant internal energy *U* is at equilibrium (i.e., maximum entropy) when the Helmholtz free energy is minimal. Substituting the definition of the internal energy and entropy (Eqs. 17 and 18, respectively), as well as the expression of *q* (Eq. 27) into Eq. 29, the Helmholtz free energy is

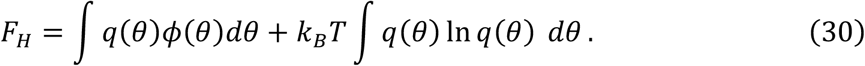

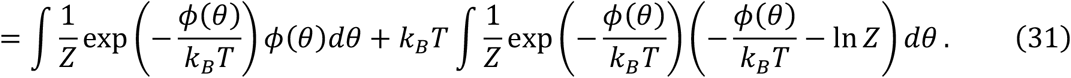

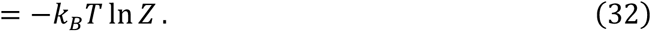

It readily follows that the log of the partition function corresponds to the negative Helmholtz free energy divided by *k_B_T*:

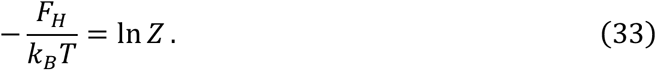

#### Free energy: A perspective from statistics

In order to link perspectives on free energy from statistical physics and (Bayesian) statistics, we assume that the system is examined at constant physical temperature *T* (which is the case if the system is connected to a large thermal reservoir). Setting the temperature so that the term *k_B_T* equals unity (normalization of temperature), we can move from a physical perspective on free energy (expressed in joules) to a statistical formulation (expressed in information units proportional to bits). This is the common convention in the statistical literature, and thereby, all quantities become unit-less information theoretic terms. Under this convention, the physical concept of free energy described above gives rises to an analogous concept of free energy in statistics when the energy function is given by the negative log joint probability −ln *p*(*y,θ|m*) (compare (Neal and Hinton, 1998):

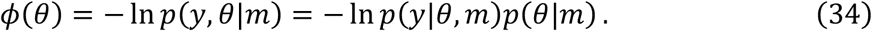

Hence, the log joint (which fully characterizes the system) takes the role of the kinetic energy in the ideal gas example above. Fig. 1 summarizes the analogies in concepts of free energy between statistical physics and statistics.

Inserting the expression for *ϕ* (Eq. 34) into Eq. 27, one can show that the equilibrium distribution of the system is the posterior distribution (i.e., the normalized joint probability):

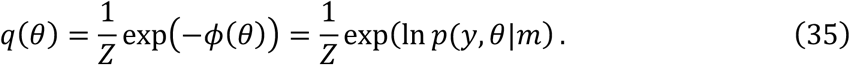

The free energy can now be written as

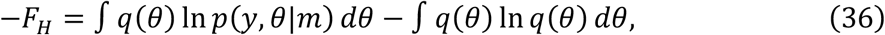

A comparison with Eq. 29 indicates that the second term in Eq. 36 should correspond to entropy; indeed, it is consistent with the definition of the Shannon entropy 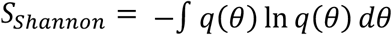. The first term, the expected log joint, corresponds to the internal energy. Finally, and most importantly for this paper, under the choice of the energy function in Eq. 34, the partition function (Eq. 28) corresponds to the normalization constant of the joint probability *p*(*y,θ|m*). By comparison with Eq. 33, we see that the negative free energy is equal to the LME:

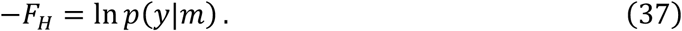

Replacing the joint in Eq. 36 by the product of likelihood and prior one can rewrite the negative free energy as

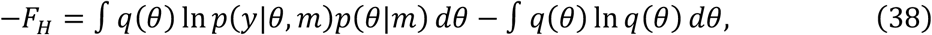

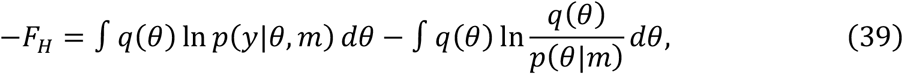

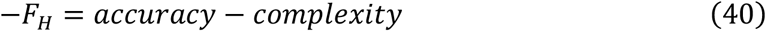

Written in this manner, the negative free energy consists of two terms that have important implications for evaluating the goodness of a model: the first term (the expected log likelihood under the posterior) represents a measure of model fit or accuracy. The second term corresponds to the Kullback-Leibler (KL) divergence between the posterior and the prior; this corresponds to the complexity of a model. Eq. 41 indicates that the negative free energy (log evidence) of a model corresponds to its balance between accuracy and complexity. We will turn to this issue in more detail below and examine variations of this perspective under TI and VB, respectively.

In the following, we will explicitly display the sign of the negative free energy for notational consistency. In order to highlight similarities with statistical physics and the concepts of energy and potential, we will continue to express the free energy as a functional of a (possibly non-normalized) log density, such that

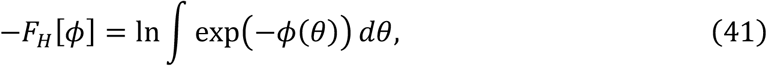

where *ϕ*(*θ*) is equivalent to an energy or potential depending on *θ*.

#### Thermodynamic integration

We now turn to the problem of computing the negative free energy. As is apparent from Eq. 41, the free energy contains an integral over all possible *θ*, which is usually prohibitively expensive to compute and thus precludes direct evaluation. One possibility to overcome this challenge is to move in small steps along a path from a known initial state to the equilibrium state and, on the way, add up changes in free energy for all steps. This idea is the basis of thermodynamic integration (TI; see Gelman and Meng, 1998) which was initially introduced in statistical physics to compute the difference in Helmholtz free energy between two states of a physical system (Kirkwood, 1935).

In Bayesian statistics, TI uses the same idea to compute the free energy (log evidence) of a model m. Here, one focuses on the transition between two “potentials” corresponding to the negative log prior density *ϕ*_0_(*θ*) = −ln *p*(*θ|m*) and the negative log joint *ϕ*(*θ*) = −1n *p*(*y|θ,m*) −ln *p*(*θ|m*). If the prior is properly normalized, i.e., 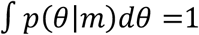, substituting *ϕ*_0_ and *ϕ* into Eq. 41 yields

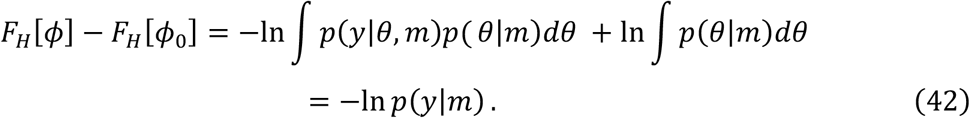

The goal is now to construct a piecewise differentiable path connecting prior and posterior and then compute the difference in free energy by integrating infinitesimal changes in the free energy along this path. A smooth transition between *F*[*ϕ*] and *F*[*ϕ*_0_] can be constructed by the power posteriors *p_β_*(*θ|y,m*) (see Eq. 50 below) which are defined by the path *ϕ_β_*:

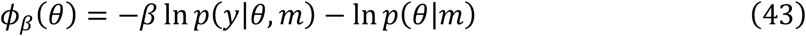

with *β* ∈ [0,1], such that *ϕ*_1_ = *ϕ*. Physically, this could be seen as studying a system under a constant potential and slowly changing a parameter *β* that controls a second potential. In the ideal gas example above, this could resemble a scenario where one studies the behavior of charged particles under the constant influence of gravity, while gradually increasing the impact of a second, for instance, electromagnetic force by increasing the electric current flowing through a coil. The equivalent perspective from statistics equates the constant potential with the negative log prior −1n *p*(*θ|m*) and the second *β* dependent potential with the negative log likelihood −ln *p*(*y|θ,m*). With growing *β*, the impact of the data increases until a balance between prior and likelihood is reached – the log joint. In the statistics literature, *β* is usually referred to as an inverse temperature because it has analogous properties to physical temperature in many aspects. We will use this terminology and comment on the analogy in more detail below.

We now turn to the construction of the path that connects the prior and the joint. It can be shown that:

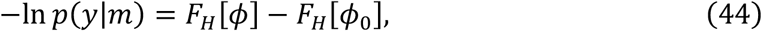

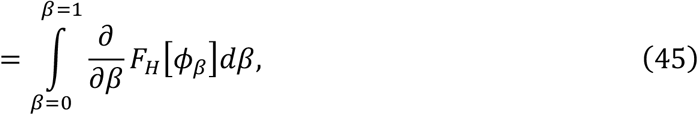

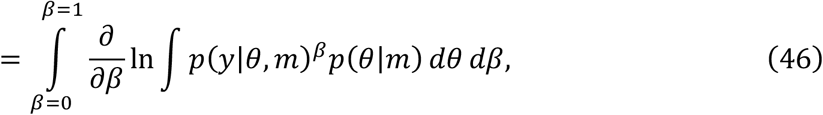

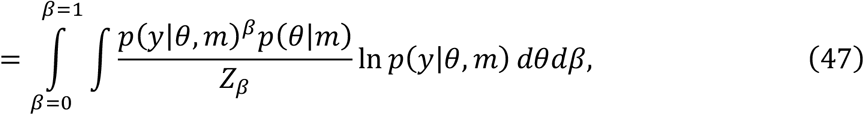

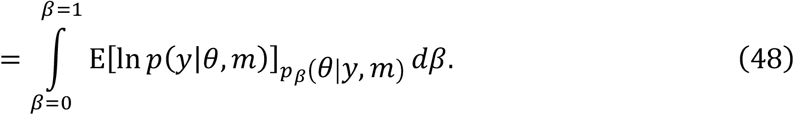

which we refer to as the basic or fundamental TI equation (Gelman and Meng, 1998). In the above equations, we have used the notation

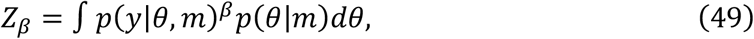

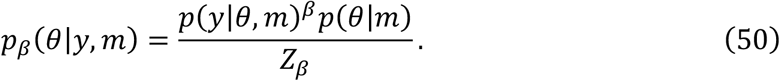

In practice, the expected value E[ln *p*(*y|θ,m*)]_*pβ*(*θ|y,m*)_ can be computed by evaluating

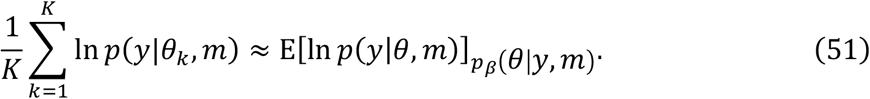

where samples *θ_k_* are drawn from the power posterior *p_β_*(*θ|y,m*). The integral over *β* in Eq. 48 can be computed through, for example, a quadrature rule using a predefined temperature schedule for *β*: 0 = β_0_ < *β*_2_ < … < *β*_*N*−1_ < *β*_*N*_ = 1. This yields

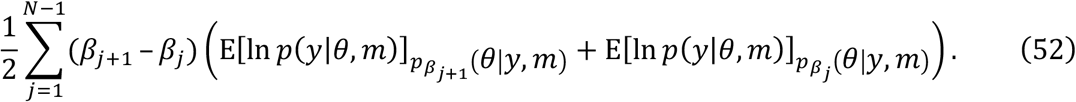

The optimal temperature schedule in terms of minimal variance of the estimator and minimal error introduced by this discretization in the context of linear models was outlined previously by Gelman and Meng (1998) and Calderhead and Girolami (2009).

To re-iterate, TI can be seen as an integral along a smooth path *l*(*β*) chosen such that *F*(*l*(0)) = *F*[*ϕ*_0_], *F*(*l*(1)) = *F*[*ϕ*].

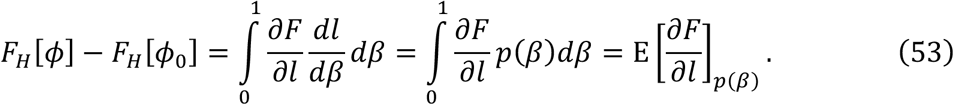

Here, we have assumed that the derivative 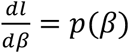 is a strictly positive function over the interval [0,1]. Thus, the selection of an optimal schedule is equivalent to the selection of an optimal importance distribution *p* (Calderhead and Girolami, 2009; Gelman and Meng, 1998). Under this perspective, TI is the expected value of the change in (negative) free energy over a set of distributions ranging from the prior to the posterior. This is in contrast to AME and HME which represent the two opposite extremes of this spectrum (Gelman and Meng, 1998; Penny and Sengupta, 2016). Intuitively, it seems plausible that TI is thus capable of providing a more faithful estimate of the log model evidence, compared to the under- and overestimation of log evidence by AME and HME, respectively.

Notably, the TI equation can also be understood in terms of the definition of the free energy by noting that the latter can be written as the sum of an expected log likelihood and a cross-entropy term (KL divergence between power posterior and prior):

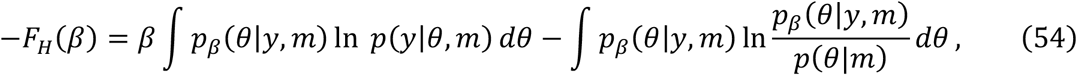

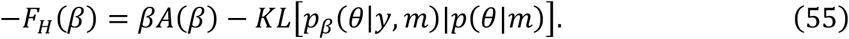

The first term, *A*(*β*) = −*∂F_H_/∂β*, is referred to as the accuracy of the model (for example Stephan et al., 2009), while the second term constitutes a complexity term. Note that Eq. 55 is typically presented in the statistical literature for the case of *β* = 1 and describes the same accuracy vs. complexity trade-off previously expressed by Eqs. 39–41, but now from the specific perspective of TI.

Fig. 2 illustrates the fundamental TI equation (Eq. 48) and shows its relation to Eq. 55. For any given *β*, the negative free energy at this position of the path −*F_H_*(*β*) can be interpreted as the signed area below the curve *A*(*β*) = −*∂F_H_/∂β*, whereas the term *β* × *A*(*β*) is the rectangular area below *A*(*β*). Eq. 55 shows that the area *βA*(*β*) + *F_H_*(*β*) is the KL divergence between the corresponding power posterior and prior.

**Figure 2:**
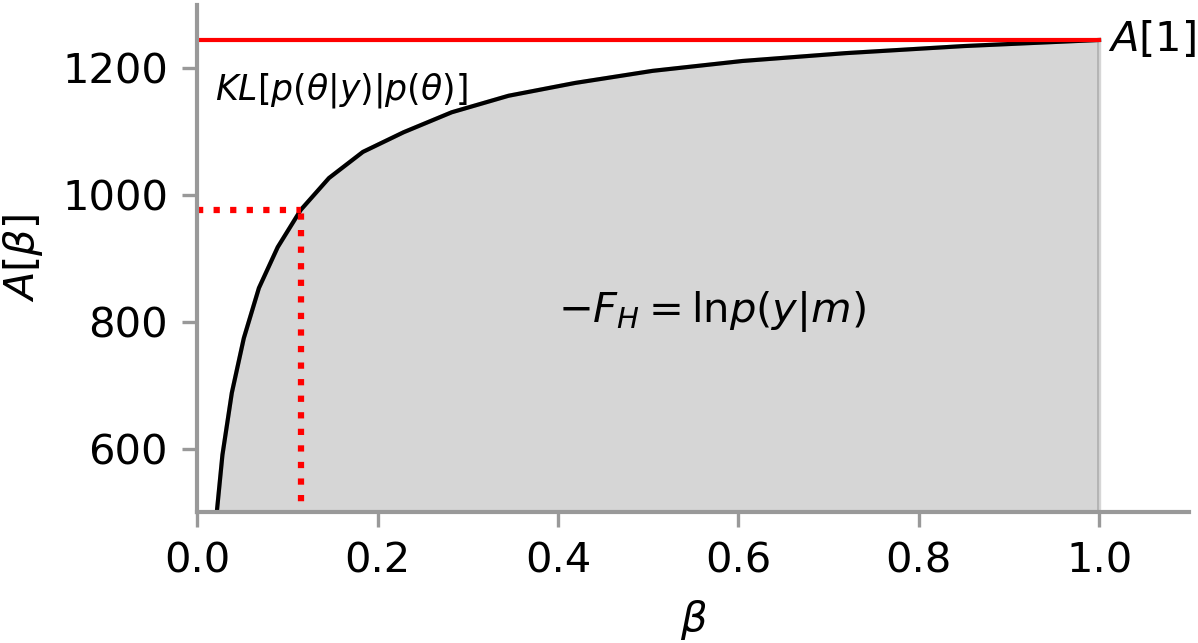
Graphical representation of the TI equation. The free energy is equal to the *signed* area below *A* = −*∂F_H_/∂β*, and thus the area *A*(1) + *F_H_* is equal to the KL divergence between posterior and prior. The same relation holds for any *β* ∈ [0,1].

This relationship holds because, for the power posteriors (Eq. 50), *A*(*β*) is a monotonically increasing function of *β*. This is due to the fact that

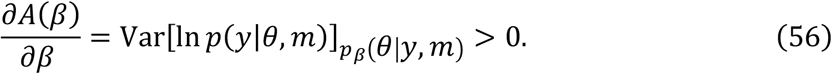

See Lartillot and Philippe (2006) for a derivation of this property. From this it follows that the negative free energy is a concave function along *β*.

The theoretical considerations highlighted above and the relation to statistical physics render TI an appealing choice for evaluating the model evidence. However, its practical utility is limited by the computational requirement of sampling from an ensemble of distributions (one for each value of *β* in Eq. 52). Arguably, it is this computational challenge that has so far prevented the use of TI in neuroimaging. Below, we present an efficient population MCMC implementation of TI that exploits parallelization and GPUs in order to overcome this bottleneck.

#### Variational Bayes

Variational Bayes (VB) is a general approach to transform intractable integrals into tractable optimization problems. Importantly, this optimization method simultaneously yields an approximation to the posterior density and a lower bound to the LME.

The fundamental equality which underlies VB is based on introducing a tractable density *q*(*θ*) to approximate the posterior *p*(*θ|y,m*).

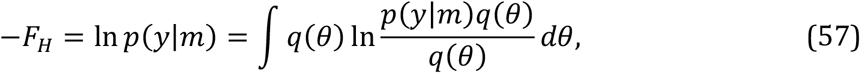

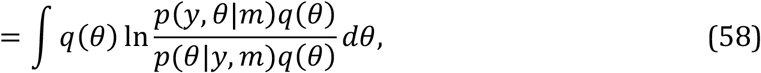

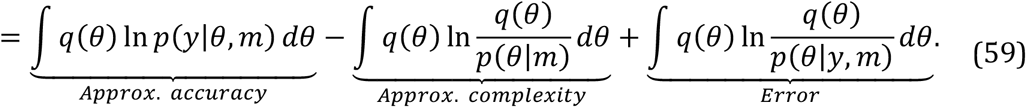

The last term in Eq. 59 is the KL divergence between the approximate density *q* and the unknown posterior density; this encodes the error or inaccuracy of the approximation. Given that the KL divergence is never negative, the first two terms in Eq. 59 represent a lower bound on the log evidence −*F_H_*, and in the following we will refer to it as the variational free energy −*F_VB_*. Eq. 59 can be rewritten as

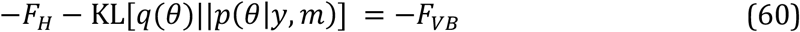

It may seem confusing that the term ‘negative free energy’ is sometimes used in the literature to denote the logarithm of the partition function *Z* itself (i.e., −*F_H_*), as we have done above, and sometimes to refer to a lower bound approximation of it (i.e., −*F_VB_*). This is because the variational free energy −*F_VB_* becomes identical to the negative free energy −*F_H_* when the approximate density *q* equals the posterior and hence their KL divergence becomes zero. In this special case

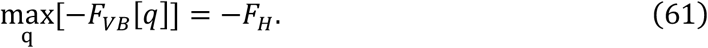

To maintain consistency in the notation, we will distinguish −*F_H_* and −*F_VB_* throughout the paper.

VB aims to reduce the KL divergence between *q* and the posterior density by maximizing the lower bound −*F_VB_* as a functional of *q*:

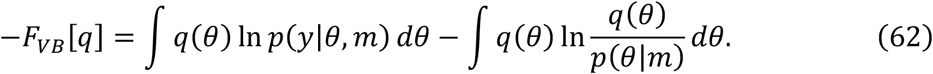

When the functional form of *q* is fixed and parametrized by a vector *η*, VB can be reformulated as an optimization method in which *η* is updated according to gradient *∂F_VB_*[*q*(*θ|η*)]/*∂η* (compare to Friston et al., 2007). Thus, the path followed by *η* can be formulated as

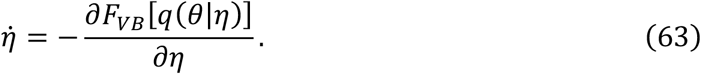

This establishes a connection between TI and VB. In the former, the path *η*(*t*) is selected a priori with the conditions that

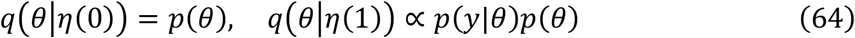

and the gradients

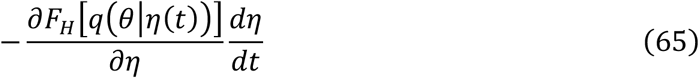

are used to numerically compute the free energy.

Different VB algorithms are defined by the particular functional form used for the approximate posterior. In the next section, we present VB under the Laplace approximation (Friston et al., 2007), as commonly used in DCM.

#### Variational Bayes under the Laplace approximation for DCM

Commonly, in order to maximize −*F_VB_*, a mean field approximation of *q* is used. In other words, the distribution *q* is assumed to factorize into different sets of parameters, each of which defines a more tractable optimization problem. In the case of DCM, *q* is assumed to have the form:

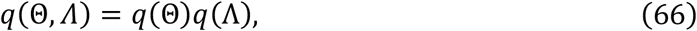

i.e., the parameters *Θ* = (*θ_c_,θ_h_,θ_g_, β*) and the hyperparameters *Λ* are assumed to be conditionally independent. The functional −*F_VB_* can be optimized iteratively with respect to *Θ* and *Λ* converging to a maximum −*F_VB_* ≤ 1n *p*(*y|m*) (Koller and Friedman, 2009). This rests on maximizing the variational energies:

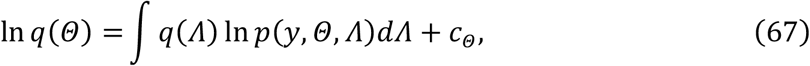

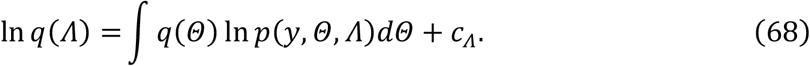

where *c_Θ_* and *c_Λ_* are constants with respect to *Θ* and *Λ*, respectively. In DCM, it is typically assumed that all terms are Gaussian (but see Raman et al., 2016 and Yao et al., 2018 who used conjugate priors for the noise terms).

Despite the mean field approximation, the integrals in Eq. 68 and 67 and cannot be solved analytically because of the nonlinearities of the forward model (Eq. 4). This problem is circumvented by approximating the log of the unnormalized posterior with a second order Taylor expansion on a local maximum (or equivalently, the unnormalized posterior is assumed to be Gaussian) and optimizing the objective function ln *p*(*y,Θ,Λ*) through gradient ascent (but see Lomakina et al., 2015 for an alternative based on Gaussian processes). This approach is called the Laplace approximation (Friston et al., 2007) and underlies other methods such as BIC (Schwarz, 1978) or when the normalization constant of an approximate, tractable posterior is directly used (Kass and Raftery, 1995). As a consequence of this approximation, the variational free energy is no longer guaranteed to represent a lower bound on the log evidence (Wipf and Nagarajan, 2009). In Supp. 2, we present a simplified version of the derivation of the VBL estimate of the free energy in (Friston et al., 2007) and an explicit expression for the accuracy term.

### Implementation

In this section, we describe the implementation for each of the estimators of the LME described above. Open source code is available in the TAPAS software package (www.translationalneuromodeling.org/tapas).

#### MCMC

TI was implemented by obtaining samples from the power posterior distributions *p_i_*(*θ|y,m*) ∝ *p*(*y|θ,m*)^*β_i_*^*p*(*θ|m*), with 10^−5^ = *β*_0_ < *β*_2_ … < *β_N_* = 1. The scaling parameter schedule obeyed a fifth order power rule as suggested by (Calderhead and Girolami, 2009). Samples from each of the chains were drawn using the Metropolis-Hastings (MH) algorithm, with a Gaussian kernel as proposal distribution for the connectivity, hemodynamic and forward model parameters (*θ_c_,θ_h_,θ_g_*). Following Shaby and Wells (2010), the covariance of the proposal distribution was modified during the burn-in phase to resemble the covariance matrix under the posterior distribution. The coefficients of the confound matrix *X*_0_ (see Eq. 4) were sampled using a Gibbs step.

The hyperparameters *Λ* were sampled using a ‘pseudo’ Gibbs step, by noting that if the prior was defined to be a Gamma distribution, its conditional posterior is again a Gamma distribution, from which samples can be easily obtained. Thus, one can replace the Gaussian prior of the log precision component by a log normal distribution and approximate it by a Gamma distribution *q*(*Λ*) with matched moments (see Raman et al., 2016), in order to obtain an analytical posterior of the form *q*(*Λ|y,Θ*)∝ *p*(*y|Θ,Λ*)*p*(*Θ*)*q*(*Λ*). This last distribution can be used to obtain samples from *Λ*. To account for this approximation, a MH step can be used as acceptance criterion for each proposed sample *Λ*\*. The corresponding ratio is

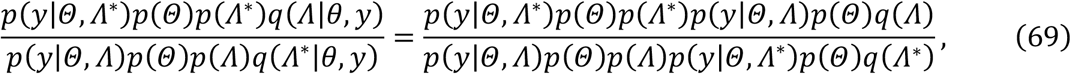

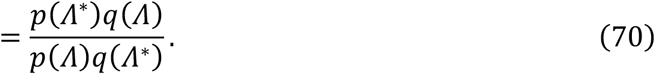

To further enhance the efficiency and convergence properties of the algorithm, we adopted a population MCMC approach in which neighboring chains were allowed to interact by means of a “swap” accept-reject (AR) step (McDowell et al., 2008; Swendsen and Wang, 1986). In brief, population MCMC defines a joint product distribution

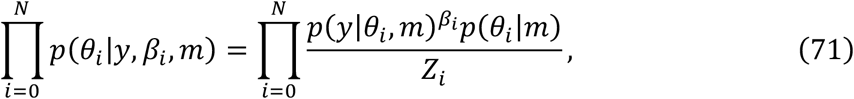

where N is the number of distributions or chains. The goal is to obtain samples from this distribution by two types of AR steps: First, local steps are used to sample parameters θ_*i*_ from *p_β_i__*(*θ_i_|y, m*). Second, samples are obtained using the swapping step in which a set of neighboring parameters *θ_i_, θ*_*i*+1_ are randomly chosen and then exchanged between chains with probability:

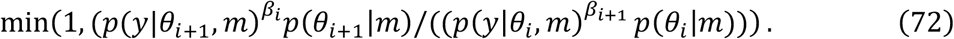

This AR step does not change the stationary distribution of any of the chains.

Population MCMC can be easily parallelized, with or without exploiting GPUs (Aponte et al., 2016) as each of the chains is independent of the rest of the ensemble. Swapping steps need to be performed serially but, assuming that the likelihood and prior functions have been already evaluated, this method increases the efficiency of the sampling scheme while only inducing negligible computational costs (for example Aponte et al., 2016; Calderhead and Girolami, 2009). Intuitively, the increase in efficiency is achieved by exploring the sampling space in a way comparable to simulated annealing, i.e., allowing some of the chains to explore the parameter space more freely by relaxing the likelihood function.

Since TI requires samples from both the prior and the posterior distribution, the same sampling algorithm can be used for computing all three sampling-based estimators (TI, AME, HME). This ensures that any observed differences between estimators are not simply due to differences in the implementation of the samplers.

We assessed the convergence of our sampling scheme using the Gelman-Rubin’s potential scale reduction factor (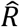 Gelman and Rubin, 1992) as diagnostic. This method tests parameter-wise convergence by comparing the variance of segments of the chains. A 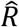 statistic below 1.1 is a commonly accepted criterion for convergence. To compute this score, the samples of the log likelihood of the first (after the burn-in phase) and last third section of each chain were compared.

#### Variational Bayes under the Laplace approximation (VBL)

The VBL algorithm used here was the implementation available in the software package SPM8 (release 5236), which employs a gradient ascent scheme to optimize the marginal distributions *q*(*Θ*) and *q*(*Λ*) (Friston et al., 2007). This algorithm was initialized at the prior mean of the parameters and hyperparameters if not stated otherwise.

To minimize any differences between our sampling-based DCM inversion approach and that of SPM, we used the same 4^th^ order Runge-Kutta scheme for integrating the DCM state equations (as required for evaluating the likelihood) in both TI and VBL.

#### Integration of the dynamical system

The computationally most intensive part of DCM is the evaluation of its likelihood function, because it requires the integration of the neuronal and hemodynamic state equations in order to predict the BOLD signal given a set of connectivity parameters. Here, we relied on the *mpdcm* toolbox (Aponte et al., 2016) which parallelizes the evaluation of the likelihood in DCM in two ways: first, the neuronal and hemodynamic states of each region are computed in parallel, i.e., the functions *f_i_* and *g_i_* are simultaneously evaluated region-wise. Second, integration can be performed for several sets of parameter values simultaneously. The integrator used here was the standard 4^th^ order Runge-Kutta explicit method. The numerical accuracy of our implementation was verified in previous work (Aponte et al., 2016).

### Simulations

#### Linear Models

In order to evaluate the accuracy of AME, HME, and TI for a situation where the ground truth is known, we first compared the estimates from a Bayesian linear regression model whose evidence can be computed analytically. These models are defined with the following prior and likelihood functions:

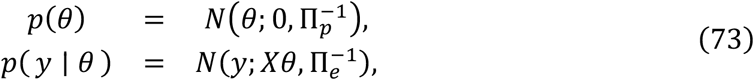

where *θ* is the [*p* × 1] vector of regression coefficients, *y* is the [*M* × 1] vector of data points, *X* is the [*M* × *p*] design matrix, and 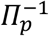 and 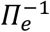 are the covariance matrices of the prior and errors, respectively. The LME is given by

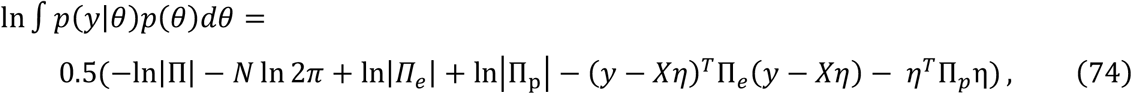

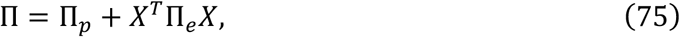

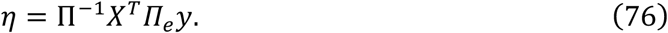

For our simulations, we chose *M* = 100, 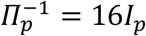 and 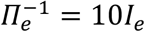, where *I_p_* and *I_e_* are the corresponding identity matrices. The design matrix was chosen to have a block structure equivalent to a design for a one-way ANOVA with *p* levels (for the values of *p* that do not exactly divide by *M*, the excess data points were assigned to the last cell). Synthetic data was generated by sampling from the generative model defined in Eq. 73.

#### DCM: Simulated data

In the first experiment, we used simulated data from 5 DCMs (linear: model 1; bilinear: models 2 to 4; nonlinear: model 5) with two inputs (*u*_1_ and *u*_2_). The DCMs are displayed in Fig. 3 and Supp. 2 and are available for download at https://www.research-collection.ethz.ch/bitstream/handle/20.500.11850/301664/simulation_dcms.zip. The parameters were chosen to maximize qualitative differences between the signals generated from them. The BOLD signal data was simulated assuming a repetition time (TR) = 2s and 720 scans per simulation. The driving inputs were entered with a sampling rate of 2.0*Hz*. Simulated time series were corrupted with Gaussian noise yielding a signal-to-noise ratio (SNR) of 1.0. Here, SNR was defined as the ratio of signal standard deviation to noise standard deviation (Welvaert and Rosseel, 2013). This means that our simulated data contained identical amounts of noise and signal, representing a relatively challenging SNR scenario.

**Figure 3:**
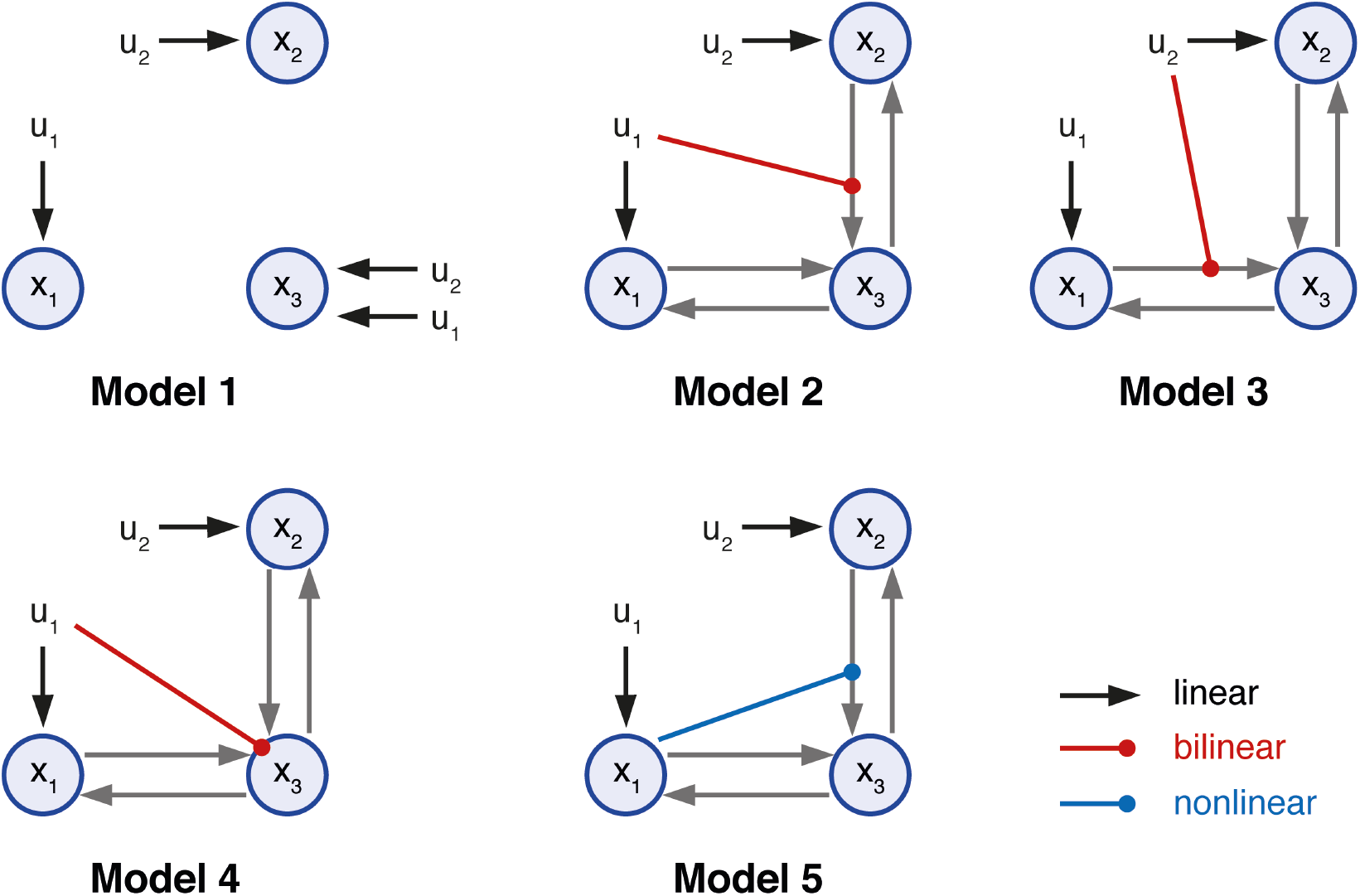
Illustration of the five simulated 3-region DCMs used for cross-model comparison. Self-connections are not displayed. The variables u_1_ and u_2_ represent two different experimental conditions or inputs. All models represented different hypotheses of how the neuronal dynamics in area x_3_ could be explained in terms of the two driving inputs and the effects of the other two regions x_1_ and x_2_. Model m_1_ can be understood as a ‘null hypothesis’ in which the activity of all the areas can be explained by the driving inputs. Models m_2_ and m_3_ correspond to two forms of bilinear effect on the forward connection of areas x_1_ and x_2_. Model nu represents the hypothesis that input u_1_ affects the self-connection of area x_3_ (not displayed). Model m_5_ represents a nonlinear interaction between regions x_1_ and x_2_. Endogenous connections are depicted by gray arrows, driving inputs by black arrows, bilinear modulations by red arrows and nonlinear modulations by blue arrows

We generated 40 different datasets with different instantiations of Gaussian noise, such that the underlying time series remained constant for each model. We then counted how often the data-generating model was assigned the largest model evidence and compared the ensuing values across the different estimators (i.e., AME, HME, TI, VBL). Notably, the absolute value of the log evidence of a given model is irrelevant for model scoring; instead, its difference to the log evidence of other models is decisive.

#### Empirical data: Attention to motion

In order to compare VBL and TI using empirical data, we used the “attention to motion” fMRI dataset (Buchel and Friston, 1997) that has been analyzed in numerous previous methodological studies (e.g., Friston et al., 2003; Marreiros et al., 2008; Penny et al., 2004a; 2004b; Stephan et al., 2008) investigated the effect of attention on motion perception; in particular, the authors examined attentional effects on the connectivity between primary visual cortex (V1), motion-sensitive visual area (V5) and posterior parietal cortex (PPC). There were four conditions (all under constant fixation): fixation only (F), presentation of stationary dots (S), passive observation of radially moving dots (N), or attention to the speed of these dots (A). Four sessions were recorded and concatenated yielding a total of 360 volumes (*T_E_* = 40*ms, TR* = 3.22*s*). Three inputs were constructed using a combination of the three conditions: *stimulus* = *S + N + A, motion = N + A, attention = A*. Driving inputs were resampled at 0.8*Hz*, requiring a total of 1440 integration steps. Further details of the experimental design and analysis can be found in Buchel and Friston (1997).

One reason for selecting this dataset is that Stephan et al. (2008) previously demonstrated that a nonlinear model had higher evidence than comparable bilinear models (Fig. 4). This case is of interest for evaluating the quality of different LME estimators, as one would expect that the introduction of nonlinearities represents a challenging case for VBL.

**Figure 4:**
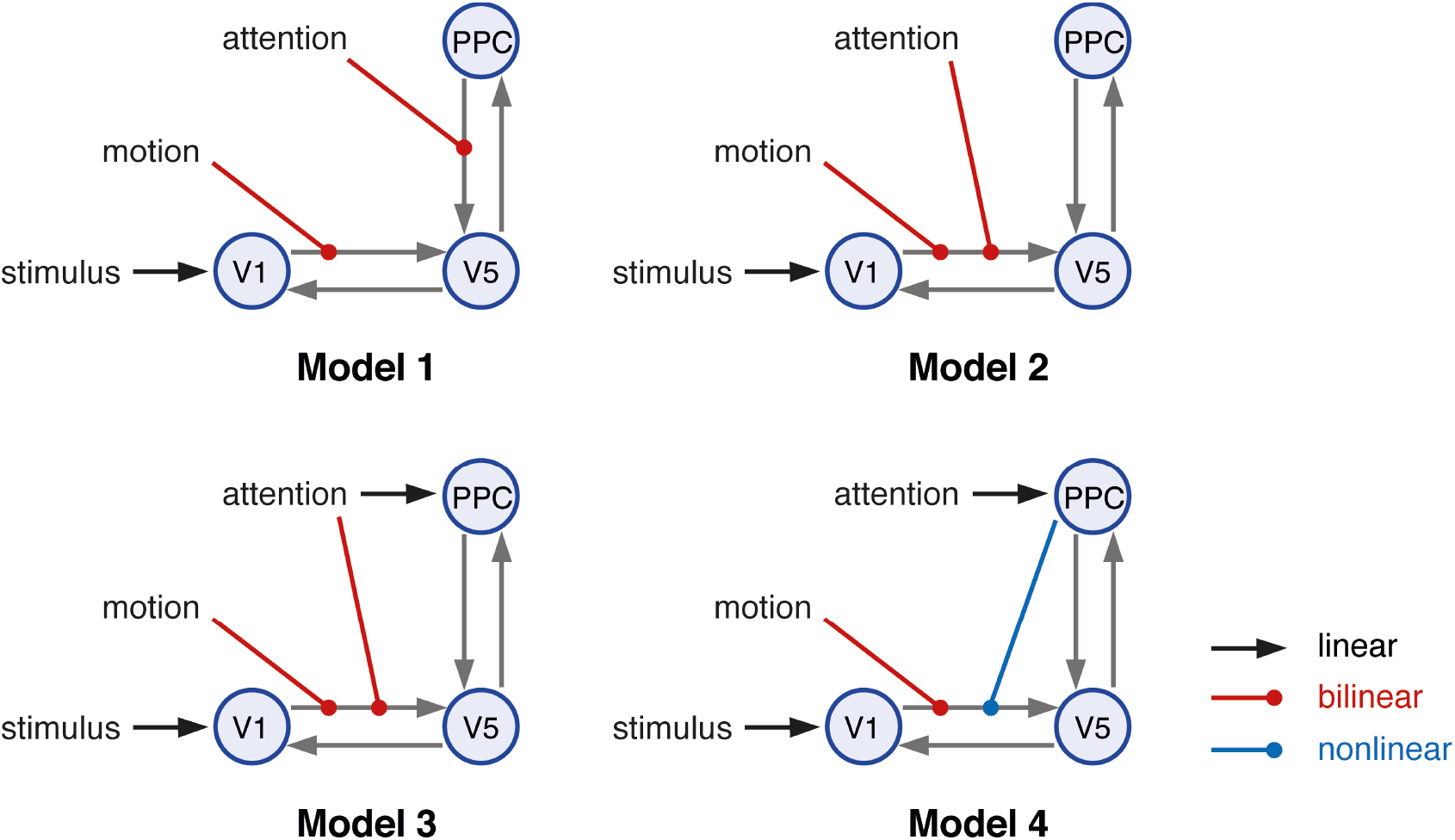
Illustration of the four models used in Stephan et al. (2008) representing different hypotheses of the putative mechanisms underlying attention-related effects in the motion-sensitive area V5. The first three models are bilinear whereas the fourth model is a nonlinear DCM. Endogenous connections are depicted by gray arrows, driving inputs by black arrows, bilinear modulations by red arrows and nonlinear modulations by blue arrows. Inhibitory self-connections are not displayed. V1: primary visual area, V5 = motion sensitive visual area, PPC: posterior parietal cortex.

#### Empirical data: Face perception

Additionally, we analyzed fMRI data from a single representative subject participating in a face perception paradigm described in detail in Frassle et al. (2016a). This dataset differs in complexity from Buchel and Friston (1997) in several ways: it consists of almost three times as many scans (940), and the DCMs contain twice the number of regions (6) and nearly three times more free parameters (shown in Figure 4, models *m*_1_, *m*_2_ and *m*_4_ possessed 28 connectivity parameters, model *m*_3_ contained 36 connectivity parameters), and a much lower SNR. This dataset is a useful candidate for evaluating sampling methods for inversion of DCMs since a previous analysis suggested possible instabilities of the VBL estimates for challenging scenarios where the number of network nodes and free parameters is high (Frassle et al., 2015).

In brief, subjects viewed either faces (F), objects (O), or scrambled (i.e., Fourier-randomized) images (S) in the left (LVF) or right visual field (RVF) in a block design, under central fixation. This study examined hemispheric lateralization in the human brain by probing intra- and interhemispheric integration in the core face perception network. The network comprised bilateral occipital face area (OFA; Puce et al., 1996) and fusiform face area (FFA; Kanwisher et al., 1997), serving as the key regions for face processing (Haxby et al., 2000), as well as left and right primary visual cortex (V1), representing the visual input regions of the network. A total of 940 scans were acquired, with TR = 1450 ms. Here, we tested the four DCMs displayed in Fig. 5. For all DCMs, non-zero entries in the endogenous connectivity (A-matrix) and driving inputs (C-matrix) were identical. Driving inputs *u*, representing the visual stimulation in either the left or right visual field, entered the contralateral V1 and were sampled at four times the frequency of the TR; thus 3760 integration steps were performed for each simulation. Forward and backward intra-hemispheric endogenous connections were assumed between V1 and OFA, and between OFA and FFA. Furthermore, reciprocal inter-hemispheric connections were set among the homotopic face-sensitive regions. Critically, models differed with regard to the experimental conditions that were allowed to perturb both intra- and interhemispheric connections, implementing different hypotheses of how hemispheric lateralization in the face perception network could arise from the functional integration within and between hemispheres. A comprehensive description of the experimental design and analysis can be found in Frassle et al. (2016b).

**Figure 5:**
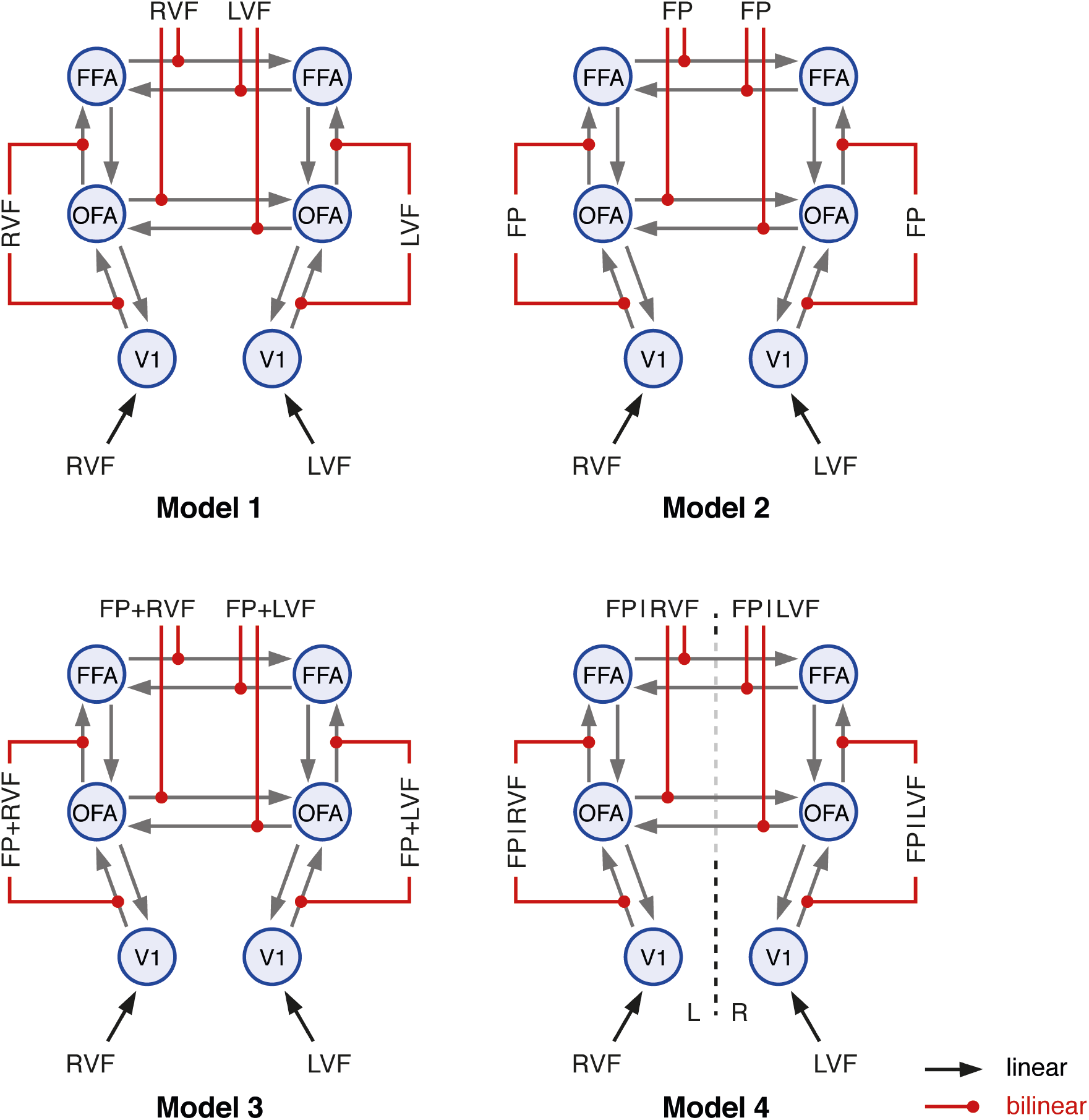
Four different models used in (Frassle et al., 2016a; 2016b) representing different hypotheses of the putative mechanisms underlying hemispheric lateralization in the face perception network. Endogenous connections are depicted by gray arrows, driving inputs by black arrows, and bilinear modulations by red arrows. Inhibitory self-connections are not displayed. V1: primary visual area, OFA: occipital face area, FFA: fusiform face area. L: left hemisphere, R: right hemisphere. LVF: left visual field, RVF: right visual field, FP: face perception. FP+RVF: Face perception and right visual field stimulation. FP+LVF: Face perception and left visual field stimulation.

## Results

### Synthetic data: linear models

In this analysis, we computed the LME of a general linear model (with a varying number *p* of regressors) using TI, AME and HME, and compared the results against the analytically computed LME.

Varying *p* from 2 to 32 in steps of 2, we repeated the data generation process 10 times. For each of these runs, values of the regression parameters *θ* were drawn from the prior, and observations *y* were generated according to the likelihood. The TI approximation to the model evidence was computed using 64 chains with a 5th order annealing schedule and 6000 samples. We then computed AME based on the samples from the prior density and HME based on samples from the posterior. Fig. 6 shows the error in the LME estimates as a function of the number of model parameters for the three approaches. Consistent with previous reports, we found that HME overestimated the LME, while AME underestimated it (Lartillot and Philippe, 2006). Only TI provided good estimates over the full range of models.

**Figure 6:**
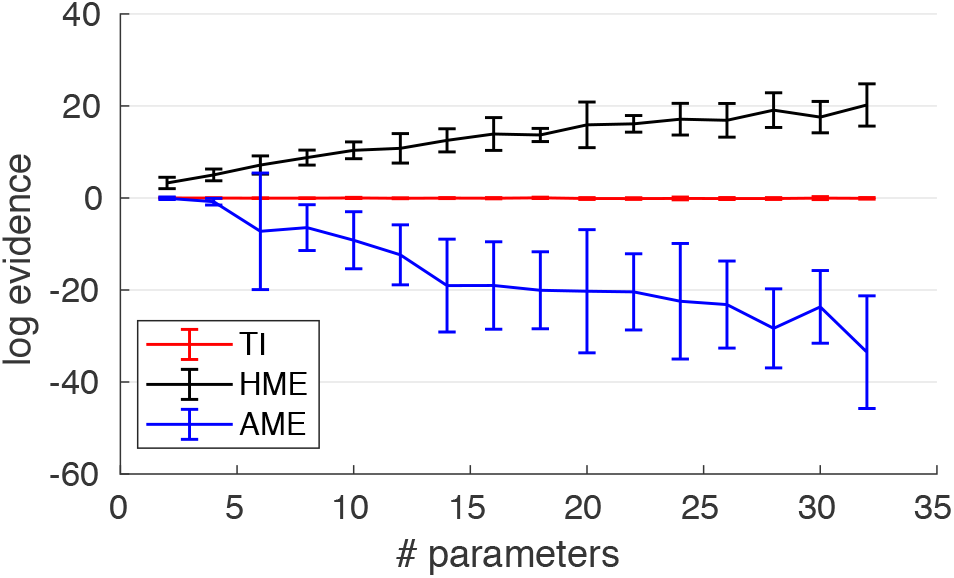
Error in estimating the log evidence of linear models for three different sampling approaches. The curves show mean and standard deviation (error bars) over ten runs at each value of p (number of GLM parameters) for thermodynamic integration (TI), posterior harmonic mean estimator (HME) and prior arithmetic mean estimator (AME).

### Synthetic data: DCM

In a pretesting phase, we found that TI generated stable estimates of the LME using 64 chains. All simulations were executed with a burn-in phase of 1 × 10^4^ samples, followed by 1 × 10^4^ kept samples. We evaluated the convergence of the MCMC algorithm by examining the samples of the log likelihood of all chains. We found that the 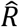 statistic was below 1.1 in all but a few instances. Estimated log model evidences are displayed in Figs. 7 and Table 1. Consistent with the linear model analysis in the previous section, the HME was always higher and the AME always lower than the TI estimate of the LME. VBL estimates were close to the TI estimate. To test for significant differences in accuracy of recovering the correct model by the different algorithms, *χ*^2^ tests were employed. TI and VBL were not significantly different (*χ*^2^ = 0.3, *p* = 0.56), but TI as well as VBL (not shown) was significantly better than AME (*χ*^2^ = 189.5, *p* < 10^−5^) and HME (*χ*^2^ = 25.4, *p* < 10^−5^).

**Figure 7:**
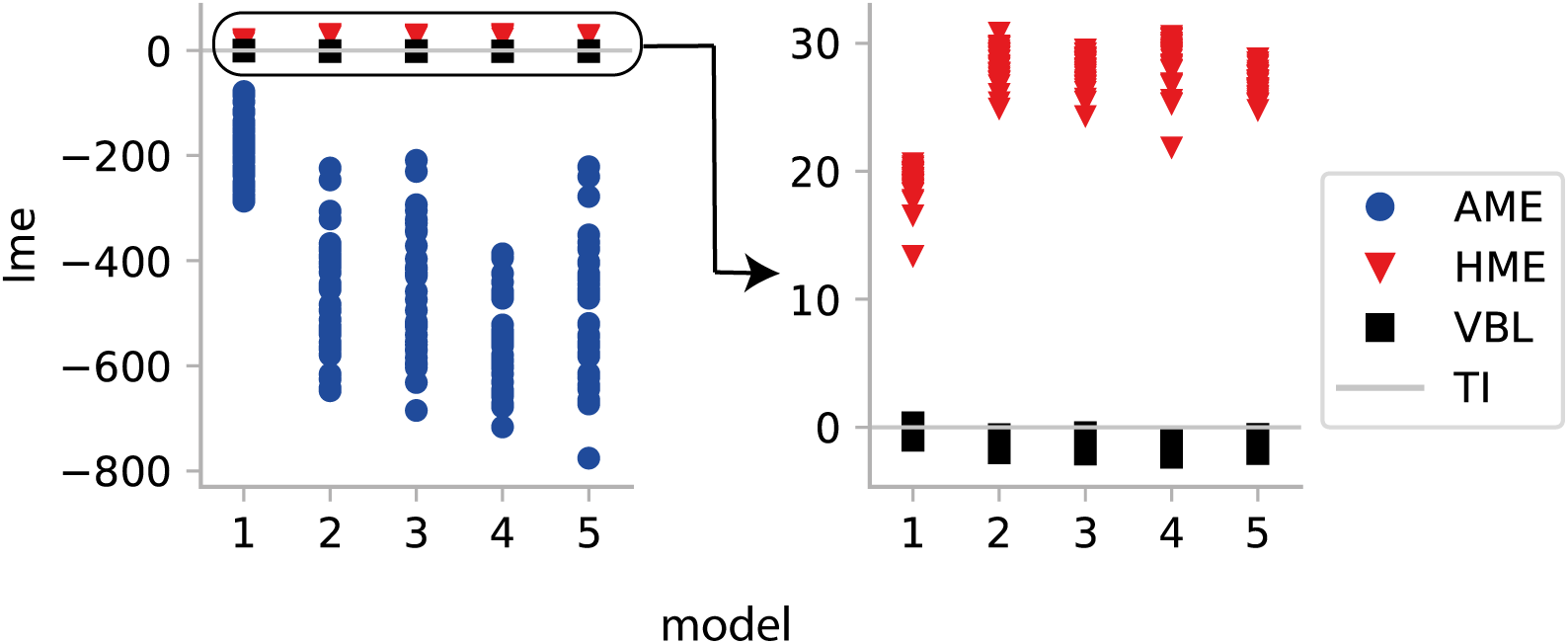
Estimated LME for all models relative to TI when inverted with the corresponding data-generating model under *SNR* = 1 for 40 different models. Right panel zooms in the left panel. Red triangles correspond to the HME, blue circles to the AME, and black squares to VBL. HME was always higher and AME always lower than the TI estimate. All LME estimates are shown after subtracting the TI-based estimate for the same model.

**Table 1:**
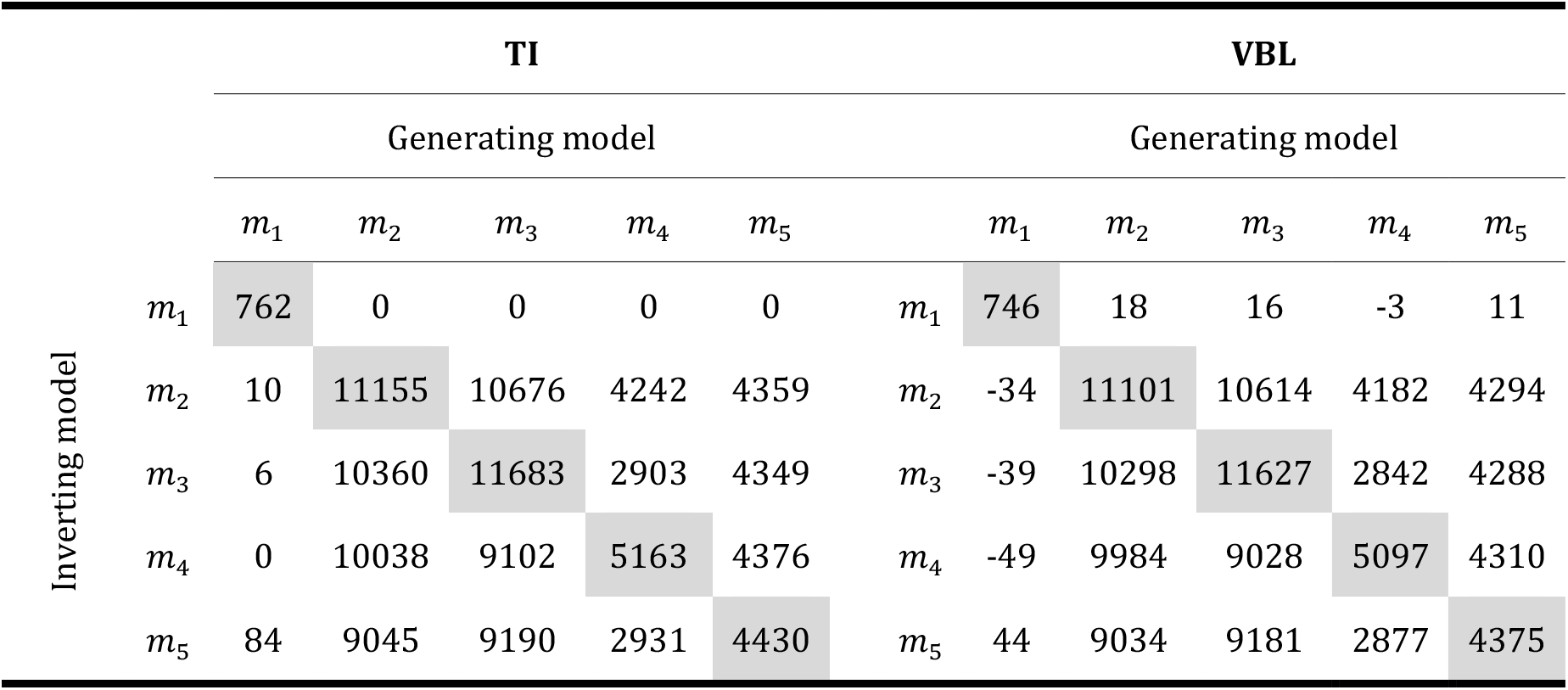
LME estimated with TI and VBL. Tables display the LME (summed across 40 simulations) of each combination of inverting and generating models. Columns have been normalized by the lowest LME: according to TI. Columns on the right and left tables share the same normalization and their absolute values can be directly compared. In most, but not in all occasion, VBL underestimated the LME compared to TI. However, for both VBL and TI the data-generating model obtained the highest LME (diagonals).

We then examined how often the data-generating model was identified correctly by model comparison, i.e., how often it showed the largest LME of all models. Of all estimators, AME failed most frequently to detect the data-generating model (Table 2). HME identified the correct model more consistently (Table 3). Both VBL and TI displayed a similar behavior (Tables 4 and 5), although model *m*_5_ was identified slightly more consistently through VBL. However, as displayed in Table 1, according to both inversion schemes, the data generating model was consistent with the model with the highest LME.

**Table 2:** Cross-model comparison results for AME in the case of synthetic data (SNR = 1). The row label indicates the data-generating model, the column index is the inferred model.

| AME: Synthetic data |  |  |  |  |  |  |
| --- | --- | --- | --- | --- | --- | --- |
| Generation |  |  |  |  |  |  |
| Inversion | | $m_1$ | $m_2$ | $m_3$ | $m_4$ | $m_5$ |
| | $m_1$ | 39 | 14 | 10 | 34 | 29 |
| | $m_2$ | 1 | 13 | 11 | | 3 |
| | $m_3$ | | 6 | 11 | 3 | |
| | $m_4$ | | 3 | 4 | 2 | 5 |
| | $m_5$ | | 4 | 4 | 1 | 3 |

**Table 3:**
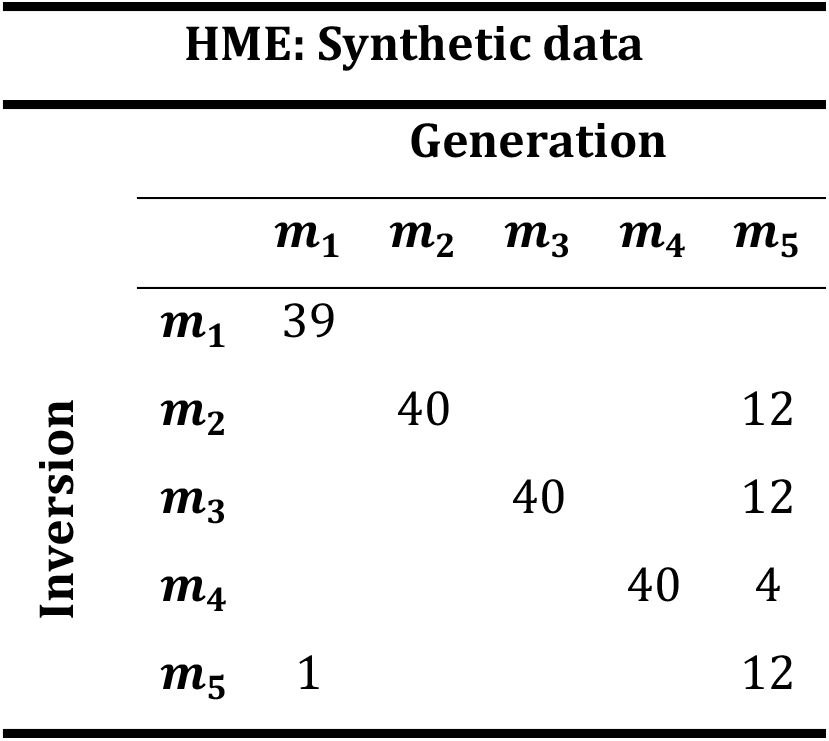
Cross-model comparison results for HME in the case of synthetic data (SNR = 1). The row label indicates the data-generating model, the column index is the inferred model.

**Table 4:**
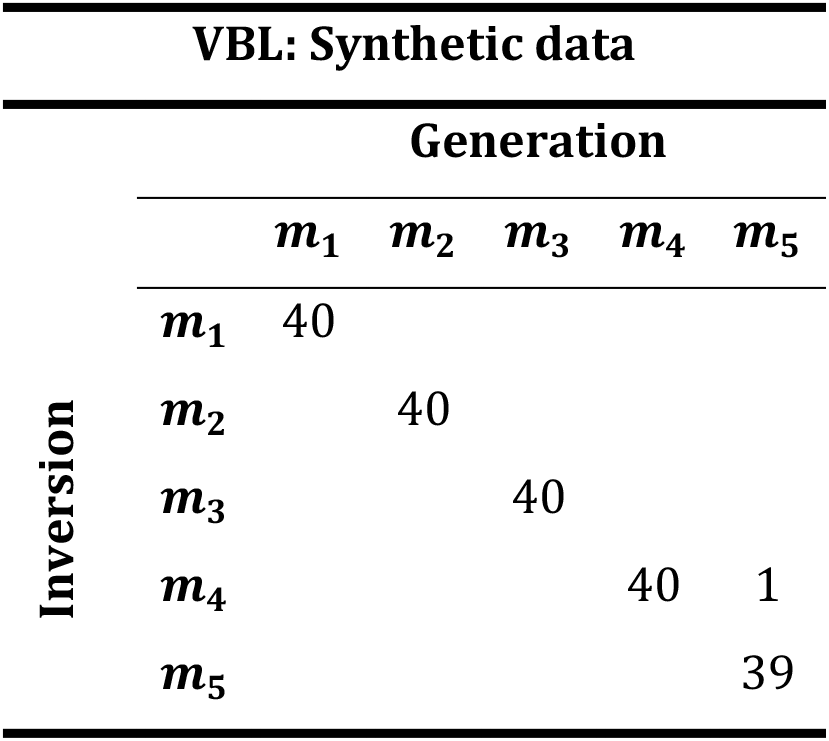
Cross-model comparison results for VBL in the case of synthetic data (SNR = 1). The row label indicates the data-generating model, whereas the column index is the inferred model.

**Table 5:**
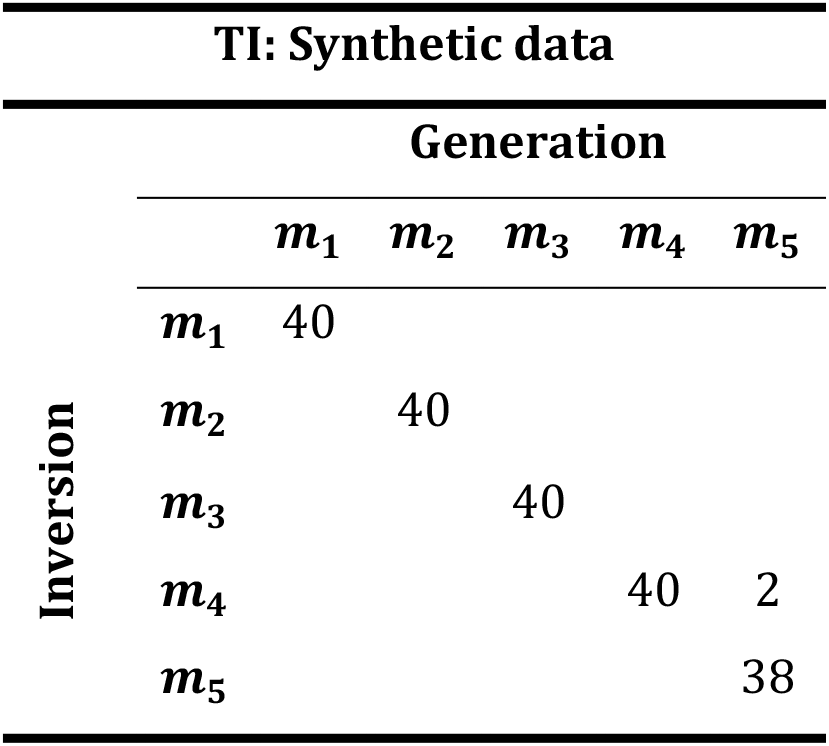
Cross-model comparison results for TI in the case of synthetic data (SNR = 1). The row label indicates the data-generating model, the column index is the inferred model.

### Empirical data

Because our previous results clearly demonstrated the inferiority of the AME and HME, in the following we limited our analysis to TI and VBL.

#### Attention to Motion

For the attention to motion dataset, 16 × 10^3^ samples were collected from 64 chains, of which 8 × 10^3^ were discarded in the burn-in phase. The convergence of the algorithm was evaluated using the 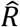 statistic of the samples of the log likelihood of each chain and model. In all but one chain, 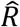 was below 1.1, indicating convergence.

Table 6 summarizes the evidence estimates obtained with TI and VBL. In comparison to the previous results by Stephan et al. (2008) Table 8, three findings are worth highlighting. First, as shown in Table 6, the VBL algorithm reproduced the ranking of models reported in Stephan et al. (2008), although an earlier version of the VBL algorithm with different prior parameters and a different integration scheme was used by Stephan et al. Moreover, our TI implementation produced the same ranking as the one obtained under VBL.

**Table 6:**
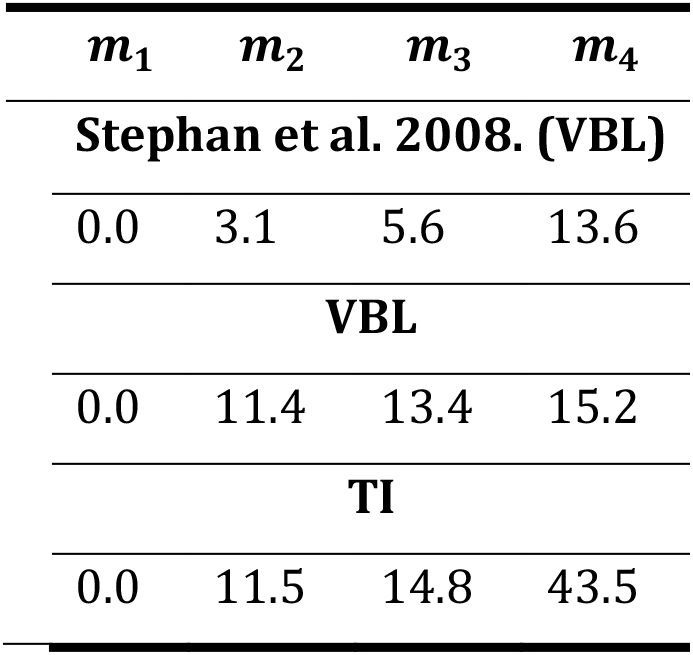
Results of model comparison, in terms of log evidence differences with respect to the worst model (m_1_), from Stephan et al. (2008), who used a different prior and integrator as in here.

Second, the difference between the VBL free energy estimates and the TI estimates varied considerably across models. To investigate this variability, we compared TI and VBL with regard to the accuracy term. The results are summarized in the lower section of Table 7. Table 7 shows that the discrepancies between VBL and TI varied across models, and the difference was particularly pronounced for the nonlinear model *m*_4_ (>40 log units).

**Table 7:** Log model evidence, accuracy and log likelihood at the MAP estimate using both TI and VBL.

| Attention to motion dataset |  |  |  |  |
| --- | --- | --- | --- | --- |
| Log model evidence |  |  |  |  |
| | $m_1$ | $m_2$ | $m_3$ | $m_4$ |
| VBL | -1790.0 | -1778.6 | -1776.6 | -1774.8 |
| TI | -1772.6 | -1761.1 | -1757.8 | -1729.1 |
| Accuracy |  |  |  |  |
| | $m_1$ | $m_2$ | $m_3$ | $m_4$ |
| VBL | -1547.6 | -1538.5 | -1531.6 | -1530.7 |
| TI | -1525.6 | -1520.2 | -1511.8 | -1483.5 |

Third, while VBL detected the most plausible model, the findings from this dataset suggest that VBL-based inversion of DCMs might not always be fully robust. In particular, the difference between the algorithms could be attributed to the VBL algorithm converging to a local extremum. To assess the differences between TI and VBL more systematically, we initialized each algorithm 10 times from different starting values that were randomly sampled from the prior density. Fig. 8 depicts the estimated model evidence and accuracy and Supp. Fig. 1 displays the predicted BOLD signal. VBL estimates of the accuracy and LME displayed much larger variance than the TI estimates. This suggests that the greater variance of the VBL estimates is due to the propensity of the gradient ascent used in VBL to converge to local maxima.

**Figure 8:**
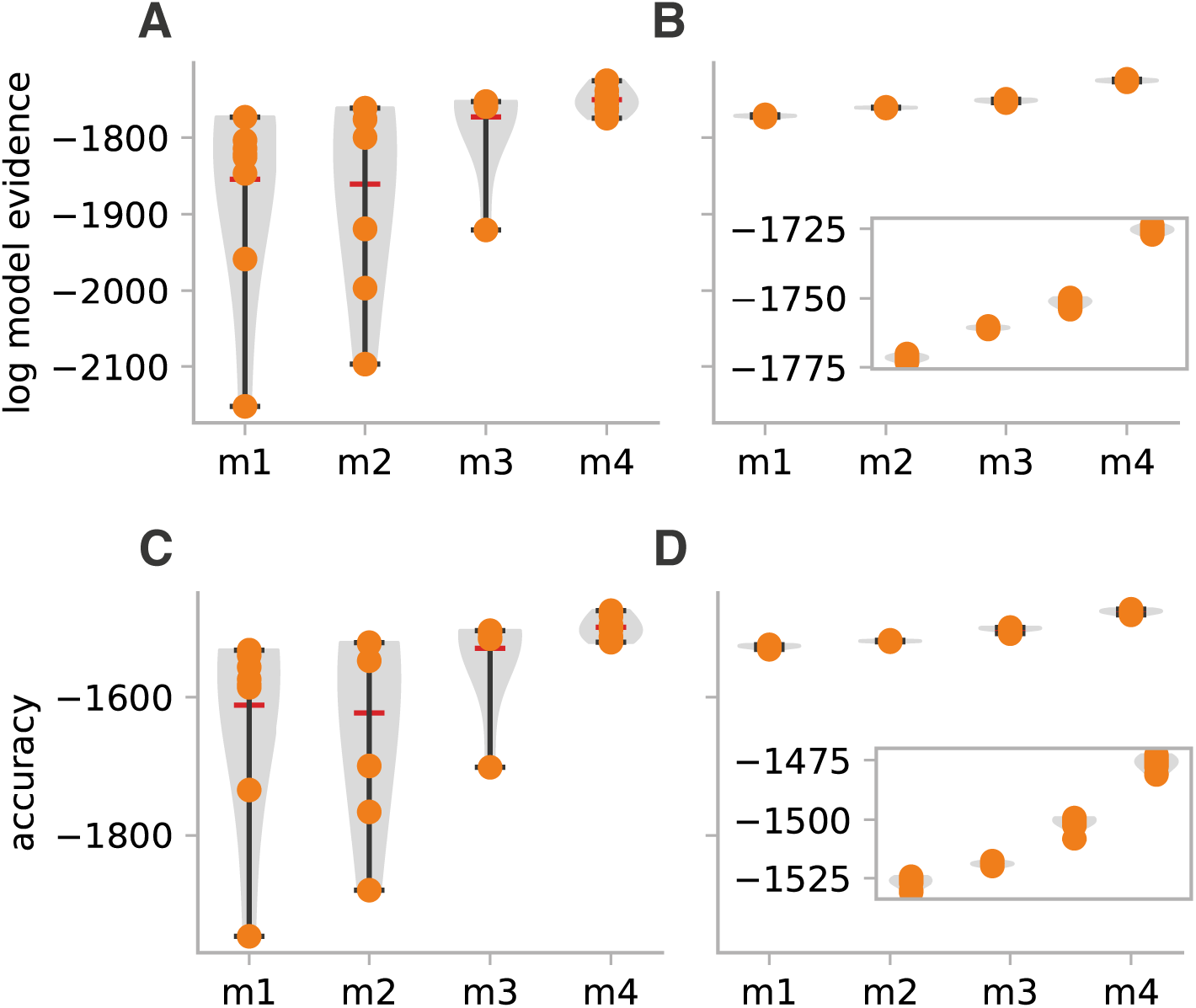
Estimates of the LME and accuracy in the attention to motion dataset after initializing VBL and TI from 10 different starting points (yellow points) drawn from the prior. The inset on the right panel zooms into the range of TI estimates. **A.** LME estimates from VBL. **B.** LME estimates from TI. **C.** LME estimates from VBL. **D.** LME estimates from TI. TI estimates show much lower variability as compared to VBL estimates.

#### Face perception

For the face perception dataset, the same number of iterations (16 × 10^3^), and discarded burn-in samples (8 × 10^3^) were used, but the number of chains was increased to 96. We found that for all but 3 chains (<1%), the 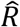 was below 1.1, indicating the convergence of the algorithm. The difference between TI and VBL estimates of the LME, accuracy and log likelihood at the MAP are shown in Table 8. Again, differences between the estimates were apparent, but more importantly there were differences in the actual ranking of the models. In particular, while VBL favored model *m*_1_, TI favored model *m*_3_. Interestingly, although the estimates of the LME showed large differences, the estimates of the accuracy were similar for the two inversion schemes.

**Table 8:** Log model evidence and accuracy estimated with TI and VBL for the four DCMs of the “face perception” dataset.

| Face perception dataset |  |  |  |  |
| --- | --- | --- | --- | --- |
| | $\mathbf{m}_1$ | $\mathbf{m}_2$ | $\mathbf{m}_3$ | $\mathbf{m}_4$ |
| Log model evidence |  |  |  |  |
| VBL | -3060.8 | -3088.4 | -3071.6 | -3065.8 |
| TI | -3051.5 | -3059.7 | -3032.9 | -3051.8 |
| Accuracy |  |  |  |  |
| VBL | -2303.6 | -2313.7 | -2279.6 | -2307.4 |
| TI | -2309.9 | -2317.9 | -2279.8 | -2303.2 |

To better understand the observed differences between the algorithms, we repeated the simulations 10 times by sampling the starting points of VBL and TI from the prior, after scaling its variance by 10^−0.5^. The scaling was used to avoid numerical instabilities encountered with the VBL algorithm. Results for the LME and accuracy under the different starting positions are shown in Fig. 9. Consistent with the attention to motion dataset, the variance of the VBL estimates was much higher than the TI estimates.

**Figure 9:**
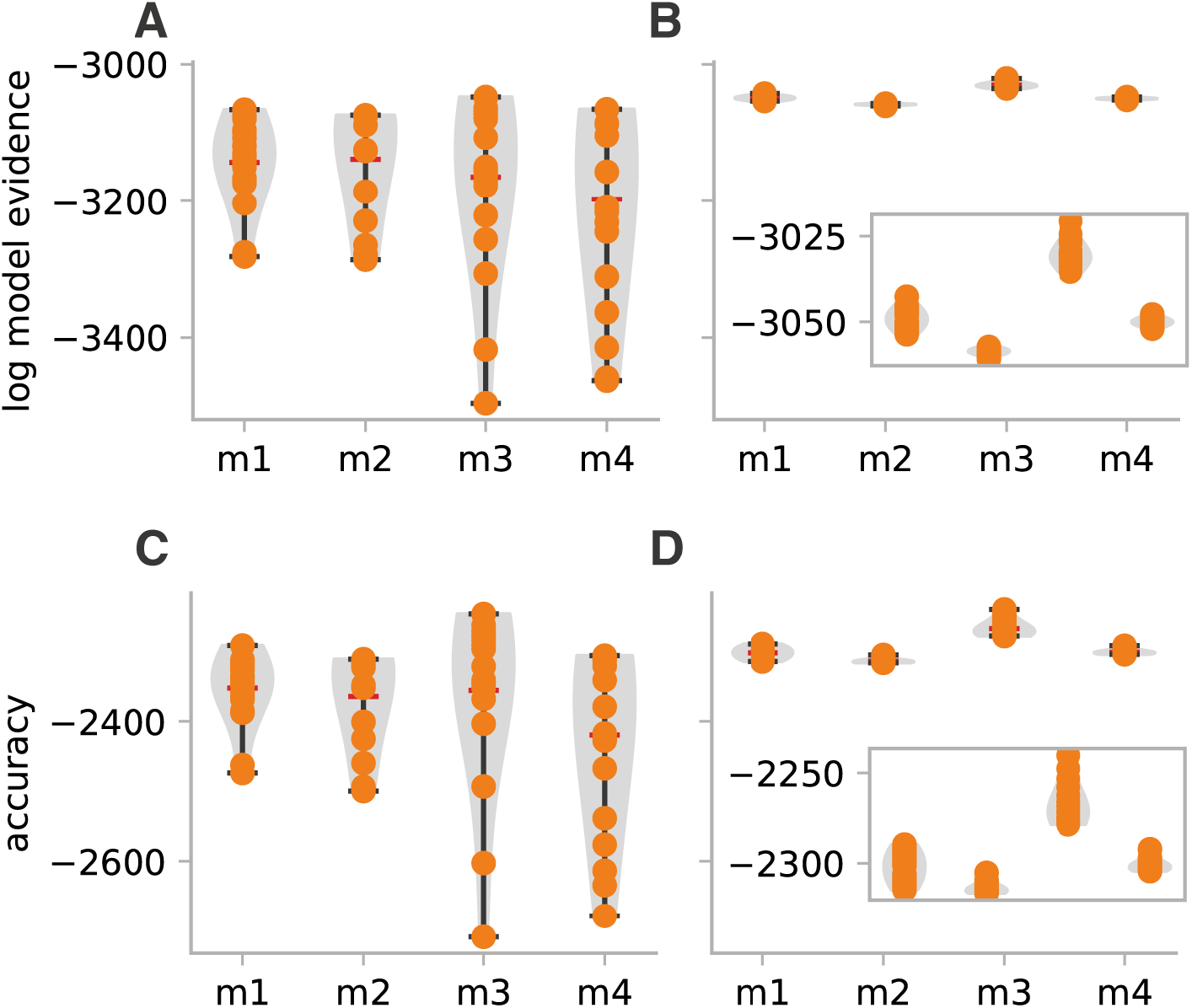
Estimates of the LME and accuracy in the face perception dataset after initializing VBL and TI from 10 different expansion points drawn from the prior. **A.** LME estimates from VBL. **B.** LME estimates from TI. **C.** LME estimates from VBL. **D.** LME estimates from TI. Clearly, TI estimates show lower variability. The inset on the right panel zooms in the TI estimates.

This was also apparent when inspecting the predicted BOLD signal time series from TI and VBL for the model with the highest score (*m*_3_) under the different initializations of the two algorithms (Sup. Fig. 2). Both VBL and TI generated qualitatively similar predictions; however, TI yielded more consistent results, suggesting that gradient ascent optimization was affected by its vulnerability to local maxima.

## Discussion

In this technical note, we have described in detail various options for approximating the model evidence of probabilistic generative models. In brief, our analyses gave three main results. First, we replicated previous findings (for example Lartillot and Philippe, 2006) that HME and AME exhibit inadequate performance and are not well-suited for estimating the LME. Having said this, variants of both estimators have been proposed that solve some of these known problems (Penny and Sengupta, 2016).

Second, TI provided robust estimates of LME, with superior performance compared to other estimators. It therefore represents a promising method for particularly challenging generative models, such as DCMs of electrophysiological data (Penny and Sengupta, 2016; Sengupta et al., 2016; 2015) or hierarchical models of DCM (Raman et al., 2016).

Third, although VBL was robust in most instances, we found evidence for variability in the estimates due to local optima in the objective function – especially for nonlinear models or challenging scenarios where the number of network nodes and free parameters is high. While its computational efficiency and relatively robust performance justify VBL as a default choice for standard applications of DCM for fMRI, sampling-based approaches like TI might become the method of choice when the robustness and validity of single-subject inference is paramount. For example, the utility of generative models for clinical applications, such as differential diagnosis based on model comparison or prediction of individual treatment responses (Stephan et al., 2017b), heavily depends on our ability to draw reliable and accurate conclusions from model-based estimates.

### Comparison between VBL and TI

Given the widespread use of VBL, its comparison to TI is of particular interest. In simulated data, both implementations yielded similar results regarding model estimates and cross-model comparison. By contrast, for the two empirical datasets tested in this paper, LME estimates differed more strongly between VBL and TI. For the “attention to motion” dataset, this was most likely due to the vulnerability of the optimization process in VBL to local optima. This was reflected by the repeated finding that different initializations (i.e., starting positions randomly sampled from the prior) led to higher variability in the VBL estimates as compared to TI.

This last observation should be interpreted in the light of several important aspects. First, the VBL algorithm used here is based on gradient optimization (Friston et al., 2007), and thus is intrinsically susceptible to local extrema. This problem can be ameliorated by initializing the optimizer from different starting points or using global optimization methods (see Lomakina et al., 2015). Second, our results are not directly comparable to previous MCMC evaluations of DCM for fMRI (Chumbley et al., 2007) as the integrator used here (Aponte et al., 2016) fully accounts for the nonlinearities of the Balloon-model. This might result in a more difficult posterior landscape than for the integrator routinely used in DCM (Friston et al., 2003), which uses a bilinear approximation. Third, the lower variability of the results obtained using MCMC reflects a trade-off between computation time and variance of the estimators. More specifically, it is not surprising that a computationally far more intensive approach like TI generates more consistent estimates as compared to the highly efficient VBL algorithm. Finally, MCMC-based methods are also susceptible to failures in convergence, and thus sampling does not constitute a silver bullet. Instead, both VBL and MCMC estimates have to be carefully examined for convergence, using over-dispersed initialization points (Gelman et al., 2003).

### Thermodynamic integration

Thermodynamic integration has received relatively little attention in the neuroimaging and cognitive science community until now (but see for example Aponte et al., 2016; 2017; Penny and Sengupta, 2016). As mentioned before, this is arguably due to the high computational burden induced by simulating from an ensemble of chains. Although the TI estimator is a computationally expensive method for computing the LME, the inherent parallel nature of this technique can be exploited to obtain estimates in reasonable time. In particular, as suggested by (Calderhead and Girolami, 2009), combining TI with population MCMC tends to increase the efficiency of the sampler and can tackle multimodal probability landscapes. This observation has been also reported by (Ballnus et al., 2017), which provided benchmark evidence that multi-chain methods can efficiently explore the posterior landscape of dynamical systems comparable to DCM, and that the computational burden is offset by increased sampling efficiency. Here, we have shown that advances in hardware allow obtaining as many as 10^5^ samples of realistic DCMs in only a few minutes. Thus, our approach provides an alternative to VBL that is available to the community as open source software (http://www.translationalneuromodeling.org/tapas/). We anticipate that further advances in sampling algorithms, specialized hardware, and software implementations will further reduce the computational time required to obtain highly accurate MCMC estimates of the model evidence. This will facilitate identification of a wide variety of nonlinear models, for which approximate methods as VBL are not adequate. Currently, the *mpdcm* toolbox (Aponte et al., 2016) supports parallelization both with Nvidia GPUs and multithreading in CPUs that does not require specialized hardware.

One of the key advantages of establishing an MCMC framework for model comparison using TI is the estimation of more complex hierarchical extensions of DCM (e.g., Raman et al., 2016). The derivation of VB equations for such hierarchical extensions is difficult (but see Friston et al., 2016; Yao et al., 2018) and may need to be revisited when models are extended or adjusted. By contrast, inference using the MCMC framework tends to be more flexible. Other important applications are cases where a multi-model posterior distribution may pose a greater challenge for inference, e.g., due to local minima. Examples beyond those discussed above include DCMs for layered fMRI (Heinzle et al., 2016) or electrophysiological signals (Kiebel et al., 2009; Penny and Sengupta, 2016) and, in particular, conductance-based DCMs (Moran et al., 2013b). Along these lines, it has been shown for DCM for EEG that gradient-based sampling-based methods outperform other, more conventional techniques, such as the Metropolis Hastings algorithm used here (Sengupta et al., 2016; 2015).

Finally. it is worth mentioning that a promising method to reduce the computational demands imposed by TI is the Widely Applicable Bayesian Information Criterion (Watanabe, 2013). This method combines the observation that, according to the intermediate value theorem there exists an optimal temperature *t*\* such that

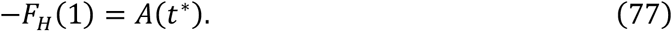

Advanced mathematical methods provide a generally valid asymptotic approximation

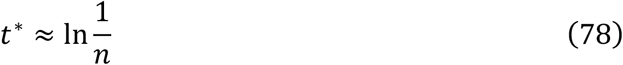

where *n* is the number of observations. This asymptotic approximation is valid even when a model does not satisfy the regularity conditions required by Laplace-based methods and traditional asymptotic approximations. This result can be particularly useful in the case of hierarchical models in case the computational burden by TI becomes prohibitive.

### Summary

In this paper, we provided a detailed tutorial-like introduction to the free energy (i) as a concept that bridges physics and statistics, and (ii) provides a foundation for TI as a principled approach to estimating the log evidence of a model. Examining the practical utility of TI in the specific context of generative models for fMRI data (DCMs), in direct comparison to VB, our analyses suggest that while VB is adequate in many instances, TI provide superior robustness, particularly in cases of nonlinear and complex models. In summary, TI-based approaches have promising potential for clinical applications where the accuracy of inference is crucial, provided sufficient computational resources are available.

## Acknowledgements

The authors acknowledge support by the René and Susanne Braginsky Foundation (KES), the Clinical Research Priority Program “Multiple Sclerosis” (KES, SR) and “Molecular Imaging” (KES) at the University of Zurich, the ETH Zurich Postdoctoral Fellowship Program and the Marie Curie Actions for People COFUND Program (SF).

## Supplementary Material

### Supp. 1

The connectivity parameters of the synthetic models used here are shown below.

#### Model 1

Model one did not include any bilinear or non-linear terms.

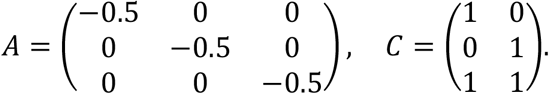

#### Model 2

Models 2 to 5 used the same A and C matrices. In addition, models 2 to 4 included one bilinear term (B matrices), and model 5 included a nonlinear term (D matrices).

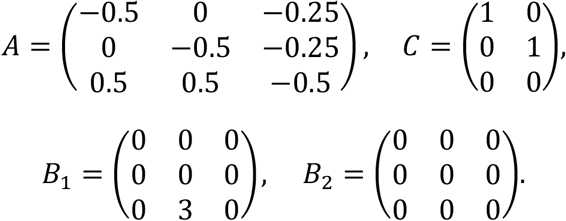

#### Model 3

Because model 3 shared the same A and C matrix with model 2, we only display the B matrices.

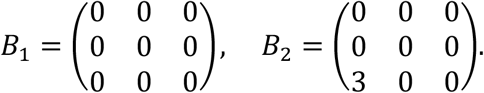

#### Model 4

Again, only the B matrices differed between models 2, 3, and 4.

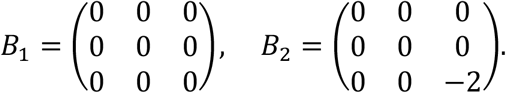

#### Model 5

Model 5 included no bilinear term but included one non-linear term.

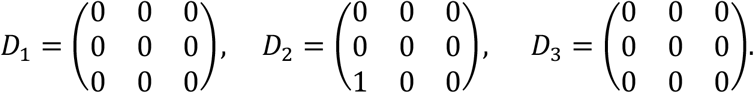

The exact data structures can be found in https://www.research-collection.ethz.ch/bitstream/handle/20.500.11850/301664/simulation_dcms.zip.

### Supp. 2

In the SPM version used here (5236), BOLD signals *y* are rescaled with respect to their *ℓ*_∞_ norm, such that

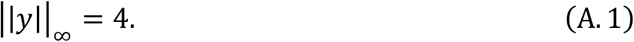

In DCM, the observation equation (see Eq. 4) can be written as

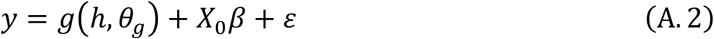

where *X*_0_ represents confounding factors. This matrix usually consists of cosine functions that account for baseline effects and low frequency components and can be imagined as implementing a model of structured noise (scanner-related fluctuations in signal intensity) that is distinct from the model’s residuals. We assume N observations such that data from a region is *y*[*t*], *t* = 0,…,*N* − 1, and the components of *X*_0_ = [*x_K_*,…,*x*_*M*−1_]^*T*^, *K* > 0 are

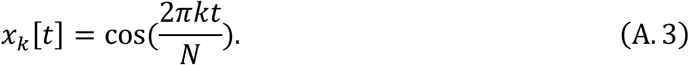

In this case, 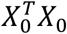 is a diagonal matrix as all base functions are orthogonal. The diagonal elements are given by

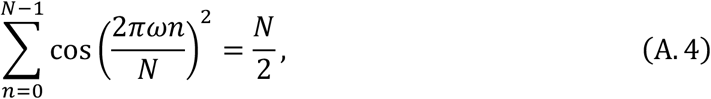

Thus,

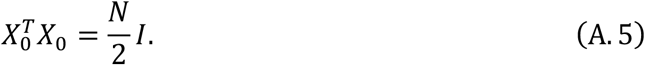

The posterior variance of the regressors conditioned on the predictions from DCM, the variance of the error 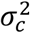, and the prior variance *σ*_0_, is

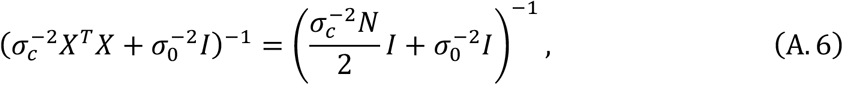

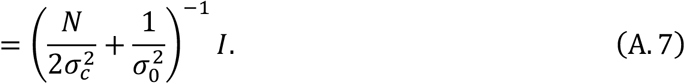

To derive the prior variance of the signal predicted by *X*_0_*β*, we note that for the predicted signal *y*:

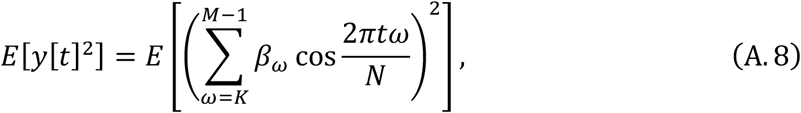

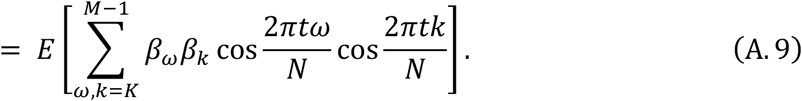

Because the coefficients are assumed to be uncorrelated and to have zero mean, it follows that

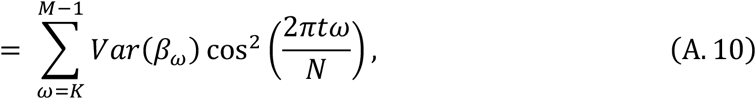

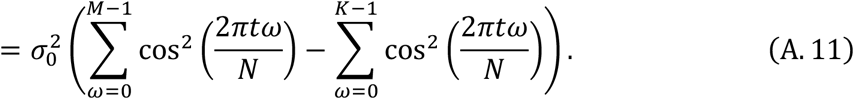

Assuming that 2*Mt/N* is an integer, it follows that

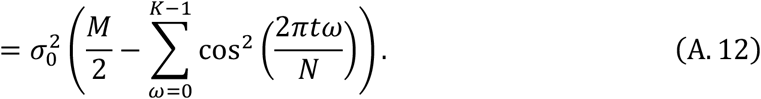

It follows that

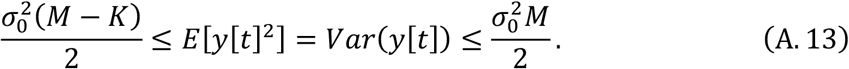

This constitutes an approximation to the prior variance of the signal. Although in the SPM implementation of DCM used here, 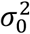 is set to 10^8^, here we use a more pragmatic value *σ*_0_ = ||*y*||_∞_ = 4. From Eq. A.12, it can be seen that this constitutes a more conservative prior variance than the SPM implementation, but still liberal enough to a priori easily account for the totality of the variance in the data.

### Supp. 3

The expression for the variational negative free energy can be derived by noting that Eq. 59 can be written as an energy term plus an entropy term

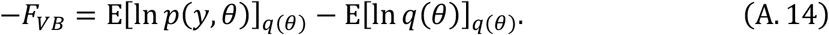

For simplicity, in the rest of this section, we collapse parameters *Θ* and hyperparameters *Λ* into a *d*-dimensional vector *θ*, assuming that a maximum has been obtained. Also, we assume that all densities are conditioned on model *m*, and make this assumption implicit. Moreover, we assume that the prior distribution of parameters *θ* is a Gaussian distribution centered at *θ*_0_ with covariance 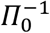.

According to the Laplace approximation, *q*(*θ*) is a Gaussian distribution with mean 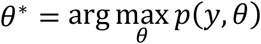 and variance

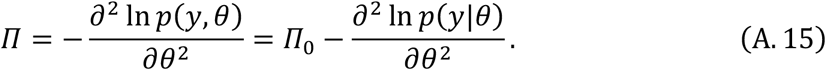

We denote the negative Hessian of the likelihood or observed Fisher information in the following as *Π_L_*.

The energy term in Eq. A.14 is approximated using the Laplace method, which yields

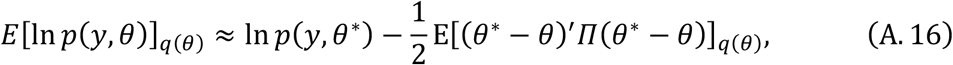

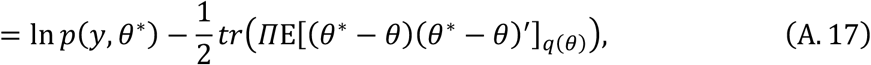

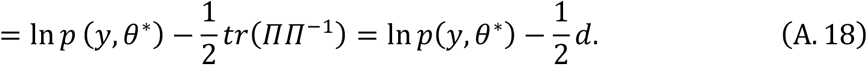

where *tr* denotes the trace operator.

The last term in Eq. A.14 is the entropy of a Gaussian distribution, which is given by:

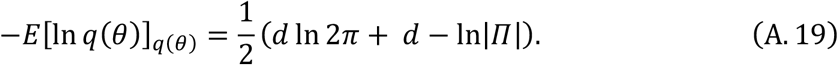

where *Π* is the precision of *q*.

Plugging Eqs. A.18 and A.19 into Eq. A.14, the variational free energy is given by

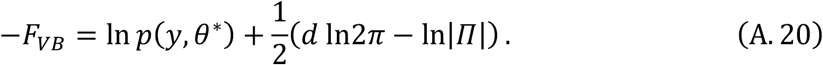

The first term on the right of Eq. A.20 can be expanded to obtain the full expression:

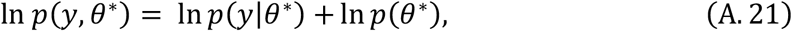

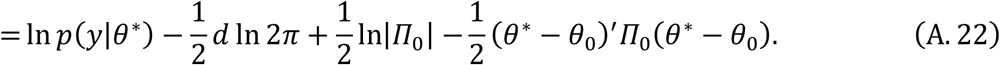

where *θ*_0_ and *Π*_0_ are the mean and precision of the prior density, respectively. By inserting Eq. into Eq. A.20, the scheme proposed by Friston et al. (2007) can be written as:

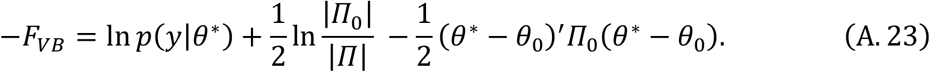

Although VBL is typically orders of magnitude faster than MCMC sampling, it exhibits several limitations: it is susceptible to (i) local extrema, (ii) violations of the distributional assumptions imposed on the posterior, (iii) violations of the conditional independence assumptions of the mean field approximation (see Daunizeau et al., 2011 for discussion), and (iv) it is only defined when the Hessian in Eq. A.15 is not singular.

Returning to our theme of connecting TI to VBL, one can write the variational negative free energy in terms of an approximate accuracy and complexity term (Eq. 59). One observes that the accuracy term can be computed as

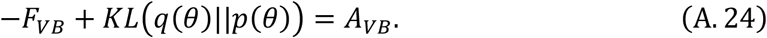

Given a Gaussian prior and posterior, the KL divergence has the following analytical form:

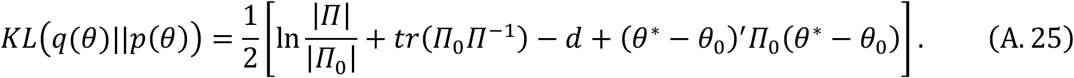

Replacing terms, we obtain

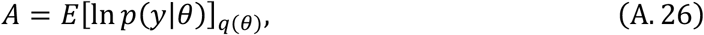

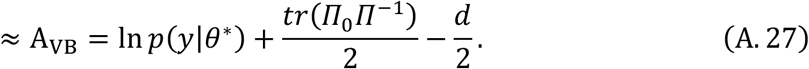

A more familiar expression for the accuracy can be derived by noting that the posterior covariance can be written as the sum of the negative Hessian of the likelihood plus the prior covariance, such that

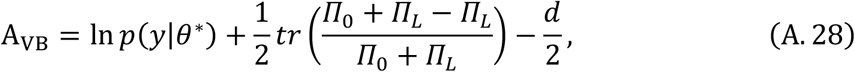

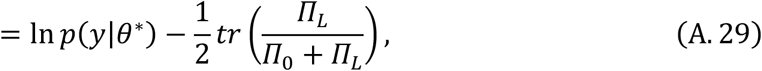

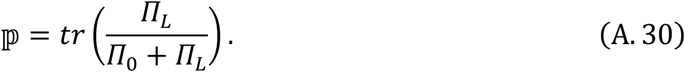

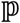 is the effective number of parameters proposed by Moody (1992) Eq. 18 and see Spiegelhalter et al. (2002) Eq. 15 and is commonly used for model selection.

### Supp. 4

**Supp. Figure 1.**
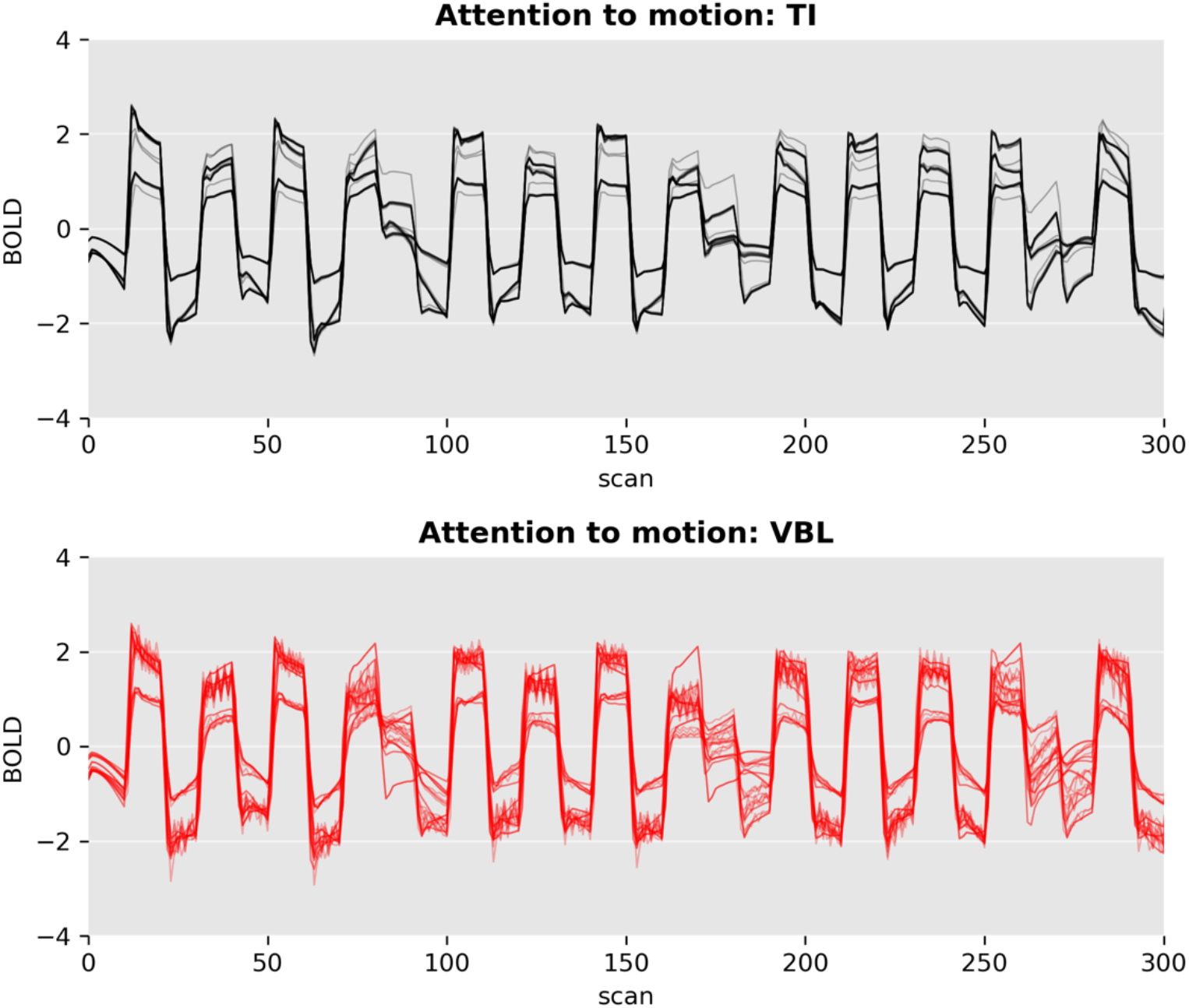
Comparison of 10 predicted BOLD signal trajectories (for the MAP estimate) of model m4 between TI and VBL for the “attention to motion” dataset from Buchel and Friston (1997). In order to obtain an unbiased impression of the variability, the predicted BOLD responses are plotted in full (i.e., including estimated confounds; compare Eq. 4). Both estimates are qualitatively similar, but VBL fits display higher variability.

### Supp. 5

**Sup. Figure 2.**
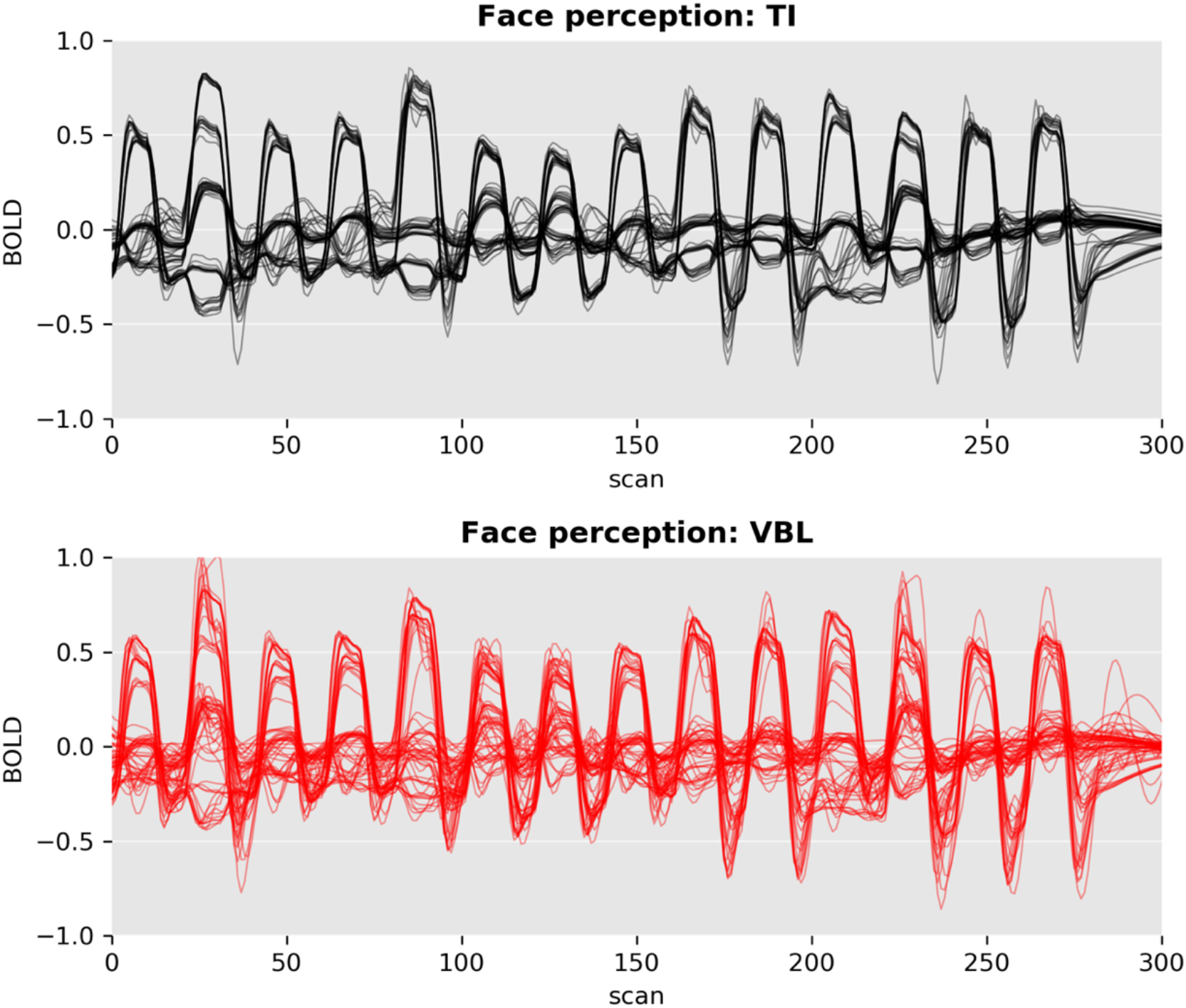
Comparison of predicted BOLD response by model *m*_3_ in Frassle et al. (2016b) for different starting points of the algorithm. Displayed are the predicted time course of 6 regions. Although both estimates are qualitatively similar, the predictions obtained under VBL display a much higher variability.

## References

Aponte, E.A., Raman, S., Sengupta, B., Penny, W.D., Stephan, K.E., Heinzle, J., 2016. mpdcm: A toolbox for massively parallel dynamic causal modeling. J. Neurosci. Methods 257, 7–16.

Aponte, E.A., Schobi, D., Stephan, K.E., Heinzle, J., 2017. The Stochastic Early Reaction, Inhibition, and late Action (SERIA) model for antisaccades. PLoS Comput Biol 13, e1005692.

Ballnus, B., Hug, S., Hatz, K., Gorlitz, L., Hasenauer, J., Theis, F.J., 2017. Comprehensive benchmarking of Markov chain Monte Carlo methods for dynamical systems. BMC Syst Biol 11, 63.

Bishop, C.M., 2006. Pattern Recognition and Machine Learning (Information Science and Statistics). Springer-Verlag New York, Inc., Secaucus, NJ, USA.

Blundell, S.J., Blundell, K.M., 2009. Concepts in thermal physics. OUP Oxford.

Buchel, C., Friston, K.J., 1997. Modulation of connectivity in visual pathways by attention: cortical interactions evaluated with structural equation modelling and fMRI. Cereb. Cortex 7, 768–778.

Buxton, R.B., Wong, E.C., Frank, L.R., 1998. Dynamics of blood flow and oxygenation changes during brain activation: the balloon model. Magn Reson Med 39, 855–864.

Calderhead, B., Girolami, M., 2009. Estimating Bayes factors via thermodynamic integration and population MCMC. Computational Statistics \& Data Analysis 53, 4028–4045. doi:http://dx.doi.org/10.1016/j.csda.2009.07.025

Chumbley, J.R., Friston, K.J., Fearn, T., Kiebel, S.J., 2007. A Metropolis-Hastings algorithm for dynamic causal models. Neuroimage 38, 478–487.

Daunizeau, J., David, O., Stephan, K.E., 2011. Dynamic causal modelling: a critical review of the biophysical and statistical foundations. Neuroimage 58, 312–322.

Frassle, S., Paulus, F.M., Krach, S., Jansen, A., 2016a. Test-retest reliability of effective connectivity in the face perception network. Hum Brain Mapp 37, 730–744.

Frassle, S., Paulus, F.M., Krach, S., Schweinberger, S.R., Stephan, K.E., Jansen, A., 2016b. Mechanisms of hemispheric lateralization: Asymmetric interhemispheric recruitment in the face perception network. Neuroimage 124, 977–988.

Frassle, S., Stephan, K.E., Friston, K.J., Steup, M., Krach, S., Paulus, F.M., Jansen, A., 2015. Test-retest reliability of dynamic causal modeling for fMRI. Neuroimage 117, 56–66.

Friston, K., Mattout, J., Trujillo-Barreto, N., Ashburner, J., Penny, W., 2007. Variational free energy and the Laplace approximation. Neuroimage 34, 220–234.

Friston, K.J., Dolan, R.J., 2010. Computational and dynamic models in neuroimaging. Neuroimage 52, 752–765.

Friston, K.J., Harrison, L., Penny, W., 2003. Dynamic causal modelling. Neuroimage 19, 1273–1302.

Friston, K.J., Litvak, V., Oswal, A., Razi, A., Stephan, K.E., van Wijk, B.C.M., Ziegler, G., Zeidman, P., 2016. Bayesian model reduction and empirical Bayes for group (DCM) studies. Neuroimage 128, 413–431.

Friston, K.J., Mechelli, A., Turner, R., Price, C.J., 2000. Nonlinear responses in fMRI: the Balloon model, Volterra kernels, and other hemodynamics. Neuroimage 12, 466–477.

Gelman, A., B, C.J., S, S.H., B, R.D., 2003. Bayesian Data Analysis. Chapman and Hall/CRC.

Gelman, A., Meng, X.L., 1998. Simulating Normalizing Constants: From Importance Sampling to Bridge Sampling to Path Sampling. Statistical Science 13, 163–185. doi:10.2307/2676756

Gelman, A., Rubin, D.B., 1992. Inference from iterative simulation using multiple sequences. Statistical Science 457–472.

Haxby, J.V., Hoffman, E.A., Gobbini, M.I., 2000. The distributed human neural system for face perception. Trends Cogn. Sci. (Regul. Ed.) 4, 223–233.

Heinzle, J., Koopmans, P.J., Ouden, den, H.E.M., Raman, S., Stephan, K.E., 2016. A hemodynamic model for layered BOLD signals. Neuroimage 125, 556–570.

Henson, R.N., Mattout, J., Phillips, C., Friston, K.J., 2009. Selecting forward models for MEG source-reconstruction using model-evidence. Neuroimage 46, 168–176.

Jaynes, E.T., 1957. Information theory and statistical mechanics. Physical review 106, 620.

Kanwisher, N., McDermott, J., Chun, M.M., 1997. The fusiform face area: a module in human extrastriate cortex specialized for face perception. J. Neurosci. 17, 4302–4311.

Kass, R.E., Raftery, A.E., 1995. Bayes factors. Journal of the american statistical association 90, 773–795.

Kiebel, S.J., Garrido, M.I., Moran, R., Chen, C.C., Friston, K.J., 2009. Dynamic causal modeling for EEG and MEG. Hum Brain Mapp 30, 1866–1876.

Kirkwood, J.G., 1935. Statistical mechanics of fluid mixtures. The Journal of Chemical Physics 3, 300–313.

Koller, D., Friedman, N., 2009. Probabilistic graphical models: principles and techniques. MIT press.

Lartillot, N., Philippe, H., 2006. Computing Bayes factors using thermodynamic integration. Syst. Biol. 55, 195–207.

Lomakina, E.I., Paliwal, S., Diaconescu, A.O., Brodersen, K.H., Aponte, E.A., Buhmann, J.M., Stephan, K.E., 2015. Inversion of hierarchical Bayesian models using Gaussian processes. Neuroimage 118, 133–145.

MacKay, D.J.C., 2003. Information Theory, Inference, and Learning Algorithms. Cambridge University Press.

MacKay, D.J.C., 2002. Information Theory, Inference \& Learning Algorithms. Cambridge University Press, New York, NY, USA.

Marreiros, A.C., Kiebel, S.J., Friston, K.J., 2008. Dynamic causal modelling for fMRI: a two-state model. Neuroimage 39, 269–278.

McDowell, J.E., Dyckman, K.A., Austin, B.P., Clementz, B.A., 2008. Neurophysiology and neuroanatomy of reflexive and volitional saccades: evidence from studies of humans. Brain Cogn 68, 255–270.

Moody, J.E., 1992. The effective number of parameters: An analysis of generalization and regularization in nonlinear learning systems, in:. Presented at the Advances in neural information processing systems, pp. 847–854.

Moran, R., Pinotsis, D.A., Friston, K., 2013a. Neural masses and fields in dynamic causal modeling. Front Comput Neurosci 7, 57.

Moran, R.J., Campo, P., Symmonds, M., Stephan, K.E., Dolan, R.J., Friston, K.J., 2013b. Free energy, precision and learning: the role of cholinergic neuromodulation. J. Neurosci. 33, 8227–8236.

Neal, R.M., Hinton, G.E., 1998. A view of the EM algorithm that justifies incremental, sparse, and other variants, in: Learning in Graphical Models. Springer, pp. 355–368.

Ortega, P.A., Braun, D.A., 2013. Thermodynamics as a theory of decision-making with information-processing costs, in:. Presented at the Proc. R. Soc. A, p. 20120683.

Penny, W., Sengupta, B., 2016. Annealed Importance Sampling for Neural Mass Models. PLoS Comput Biol 12, e1004797.

Penny, W.D., 2012. Comparing dynamic causal models using AIC, BIC and free energy. Neuroimage 59, 319–330.

Penny, W.D., Stephan, K.E., Daunizeau, J., Rosa, M.J., Friston, K.J., Schofield, T.M., Leff, A.P., 2010. Comparing families of dynamic causal models. PLoS Comput Biol 6, e1000709.

Penny, W.D., Stephan, K.E., Mechelli, A., Friston, K.J., 2004a. Comparing dynamic causal models. Neuroimage 22, 1157–1172.

Penny, W.D., Stephan, K.E., Mechelli, A., Friston, K.J., 2004b. Modelling functional integration: a comparison of structural equation and dynamic causal models. Neuroimage 23 Suppl 1, S264–274.

Puce, A., Allison, T., Asgari, M., Gore, J.C., McCarthy, G., 1996. Differential sensitivity of human visual cortex to faces, letterstrings, and textures: a functional magnetic resonance imaging study. J. Neurosci. 16, 5205–5215.

Raftery, A.E., Newton, M.A., Satagopan, J.M., Krivitsky, P.N., 2006. Estimating the integrated likelihood via posterior simulation using the harmonic mean identity.

Raman, S., Deserno, L., Schlagenhauf, F., Stephan, K.E., 2016. A hierarchical model for integrating unsupervised generative embedding and empirical Bayes. J. Neurosci. Methods 269, 6–20.

Rigoux, L., Stephan, K.E., Friston, K.J., Daunizeau, J., 2014. Bayesian model selection for group studies - revisited. Neuroimage 84, 971–985.

Robert, C., Casella, G., 2013. Monte Carlo statistical methods. Springer Science \& Business Media.

Schwarz, G., 1978. Estimating the dimension of a model. The annals of statistics 6, 461–464.

Sengupta, B., Friston, K.J., Penny, W.D., 2016. Gradient-based MCMC samplers for dynamic causal modelling. Neuroimage 125, 1107–1118.

Sengupta, B., Friston, K.J., Penny, W.D., 2015. Gradient-free MCMC methods for dynamic causal modelling. Neuroimage 112, 375–381.

Shaby, B., Wells, M.T., 2010. Exploring an adaptative Metropolis algorithm. Department of statistical science. Duke Universitiy, Durham, NC, USA.

Spiegelhalter, D.J., Best, N.G., Carlin, B.P., Van Der Linde, A., 2002. Bayesian measures of model complexity and fit. Journal of the Royal Statistical Society: Series B (Statistical Methodology) 64, 583–639.

Stephan, K.E., Iglesias, S., Heinzle, J., Diaconescu, A.O., 2015. Translational Perspectives for Computational Neuroimaging. Neuron 87, 716–732.

Stephan, K.E., Kasper, L., Harrison, L.M., Daunizeau, J., Ouden, den, H.E., Breakspear, M., Friston, K.J., 2008. Nonlinear dynamic causal models for fMRI. Neuroimage 42, 649–662.

Stephan, K.E., Penny, W.D., Daunizeau, J., Moran, R.J., Friston, K.J., 2009. Bayesian model selection for group studies. Neuroimage 46, 1004–1017.

Stephan, K.E., Schlagenhauf, F., Huys, Q.J., Raman, S., Aponte, E.A., Brodersen, K.H., Rigoux, L., Moran, R.J., Daunizeau, J., Dolan, R.J., Friston, K.J., Heinz, A., 2017a. Computational neuroimaging strategies for single patient predictions. Neuroimage 145, 180–199.

Stephan, K.E., Schlagenhauf, F., Huys, Q.J., Raman, S., Aponte, E.A., Brodersen, K.H., Rigoux, L., Moran, R.J., Daunizeau, J., Dolan, R.J., Friston, K.J., Heinz, A., 2017b. Computational neuroimaging strategies for single patient predictions. Neuroimage 145, 180–199.

Stephan, K.E., Weiskopf, N., Drysdale, P.M., Robinson, P.A., Friston, K.J., 2007. Comparing hemodynamic models with DCM. Neuroimage 38, 387–401.

Swendsen, R., Wang, J.-S., 1986. Replica Monte Carlo Simulation of Spin-Glasses. Phys. Rev. Lett. 57, 2607–2609.

Trujillo-Barreto, N.J., Aubert-Vazquez, E., Valdes-Sosa, P.A., 2004. Bayesian model averaging in EEG/MEG imaging. Neuroimage 21, 1300–1319.

Vyshemirsky, V., Girolami, M.A., 2008. Bayesian ranking of biochemical system models. Bioinformatics 24, 833–839.

Watanabe, S., 2013. A widely applicable Bayesian information criterion. Journal of Machine Learning Research 14, 867–897.

Welvaert, M., Rosseel, Y., 2013. On the definition of signal-to-noise ratio and contrast-to-noise ratio for FMRI data. PLoS ONE 8, e77089.

Wipf, D., Nagarajan, S., 2009. A unified Bayesian framework for MEG/EEG source imaging. Neuroimage 44, 947–966.

Wolpert, R.L., Schmidler, S.C., 2012. α-Stable limit laws for harmonic mean estimators of marginal likelihoods. Statistica Sinica 1233–1251.

Yao, Y., Raman, S.S., Schiek, M., Leff, A., Frassle, S., Stephan, K.E., 2018. Variational Bayesian inversion for hierarchical unsupervised generative embedding (HUGE). Neuroimage 179, 604–619.

